# Comprehensive and Comparative Transcriptional Profiling of the Cell Wall Stress Response in *Bacillus subtilis*

**DOI:** 10.1101/2023.02.03.526509

**Authors:** Qian Zhang, Charlene Cornilleau, Raphael R. Müller, Doreen Meier, Pierre Flores, Cyprien Guérin, Diana Wolf, Vincent Fromion, Rut Carballido-Lopez, Thorsten Mascher

**Author notes:** These two authors contributed equally to this work (Qian Zhang, Charlene Cornilleau).

## Abstract

The bacterial cell wall (CW) is an essential protective barrier and the frontline of cellular interactions with the environment and also a target for numerous antimicrobial agents. Accordingly, its integrity and homeostasis are closely monitored and rapid adaptive responses by transcriptional reprogramming induce appropriate counter-measures against perturbations. Here, we report a comprehensive and comparative transcriptional profiling of the primary cell envelope stress responses (CESR), based on combining RNAseq and high-resolution tiling array studies of the Gram-positive model bacterium *Bacillus subtilis* exposed to a range of antimicrobial compounds that interfere with cytoplasmic, membrane-coupled or extracellular steps of peptidoglycan (PG) biosynthesis. It revealed the complexity of the CESR of *B. subtilis* and unraveled the contribution of extracytoplasmic function sigma factors (ECFs) and two-component signal transduction systems (TCSs) to protect the cell envelope. While membrane-anchored steps are tightly controlled, early cytoplasmic and late extracellular steps of PG biosynthesis are hardly monitored at all. The ECF σ factors σ^W^ and particularly σ^M^ provide a general CESR, while σ^V^ is almost exclusively induced by lysozyme, against which it provides specific resistance. Remarkably, σ^X^ was slightly repressed by most antibiotics, pointing towards a role in envelope homeostasis rather than CESR. It shares this role with the WalRK TCS, which balances CW growth with controlled autolysis. In contrast, all remaining TCSs are envelope stress-inducible systems. LiaRS is induced by a wide range of PG synthesis inhibitors, while the three paralogous systems BceAB, PsdRS and ApeRS are more compound-specific detoxification modules. Induction of the CssRS TCS by all antibiotics interfering with membrane-anchored steps of PG biosynthesis points towards a physiological link between CESR and secretion stress. Based on the expression signatures, a suite of CESR-specific *B. subtilis* whole cell biosensors were developed and carefully evaluated. This is the first comprehensive transcriptomic study focusing exclusively on the primary effects of envelope perturbances that shall provide a reference point for future studies on Gram-positive CESR.

## Introduction

The bacterial cell envelope separates and protects the cell from its environment. It serves as a molecular sieve, a diffusion barrier, and a communication interface and counteracts the high internal osmotic pressure [1, 2]. In Gram-positive bacteria, it consists of the cytoplasmic membrane surrounded by a thick CW, primarily composed of two biopolymers, the peptidoglycan (PG) and the anionic wall teichoic acids. PG forms a three-dimensional network that maintains cell shape and provides physical integrity by counteracting the very high internal osmotic pressure of bacterial cells.

### The bacterial cell wall as shield and target

PG is made of glycan chains of alternating N-acetyl- glucosamine (GlcNAc) and N-acetyl-muramic acid (MurNAc), cross-linked by stem peptides linked to MurNAc and synthesized as pentapeptides. It is assembled in three major steps that are confined to different cellular compartments (Fig. 1): (i) the cytoplasmic assembly of soluble UDP-GlcNAc and UDP-MurNAc-pentapeptide, (ii) the membrane-associated formation of the lipid II intermediate and its translocation across the cytoplasmic membrane, referred to as the lipid II-cycle, and (iii) the incorporation and crosslinking of the GlcNAc-MurNAc-pentapeptide building block into the established PG network by transglycosylation (TG) and transpeptidation (TP) reactions [3]. After releasing the building blocks, the lipid carrier (undecaprenyl phosphate) flips back to the inner leaflet of the cell membrane and is then recycled for the next round of translocation (Fig. 1). Because of its essential function, the CW represents an attractive target for antimicrobial compounds, especially since the PG layer is a uniquely bacterial structure not found in eukaryotes. Thus, PG biosynthesis inhibitors, such as the β-lactams, display low target-related side effects and are still the most widely used antibacterial drugs worldwide [4]. Basically every step of the essential PG biosynthesis pathway is targeted by antibiotics [5] (Fig. 1). Fosfomycin and D-cycloserine are inhibitors of the cytoplasmic steps of precursor biosynthesis. Fosfomycin blocks the first committed step, the formation of UDP-MurNAc from UDP-GlcNAc by inhibiting the catalytic enzyme MurA [6]. D-cycloserine inhibits both the D-alanine racemase Alr and the D-Ala-D-Ala ligase DdlB that produce UDP-MurNAc-pentapeptide [7–9]. A plethora of antibiotics interfere with the membrane-anchored steps of PG biosynthesis: tunicamycin at high concentrations (>10 µg/ml) blocks MraY activity in addition to the target of surface glycopolymers, thereby inhibiting the formation of lipid I (MurNAc-pentapeptide-UPP) from UDP-MurNAc-pentapeptide [10–14], while bacitracin binds to undecaprenyl pyrophosphate, thereby preventing its dephosphorylation and hence recycling of the lipid carrier [15–17]. Vancomycin is a glycopeptide antibiotic that blocks glycan polymerization and cross-linking by binding to the D-alanyl-D-alanine dipeptide terminus of newly externalized lipid II [18]. Finally, moenomycin inhibits the TG reaction of PBPs [19, 20], while the β-lactams (e.g. Penicillin G) interfere with their TP reaction [21].

**Fig. 1.**
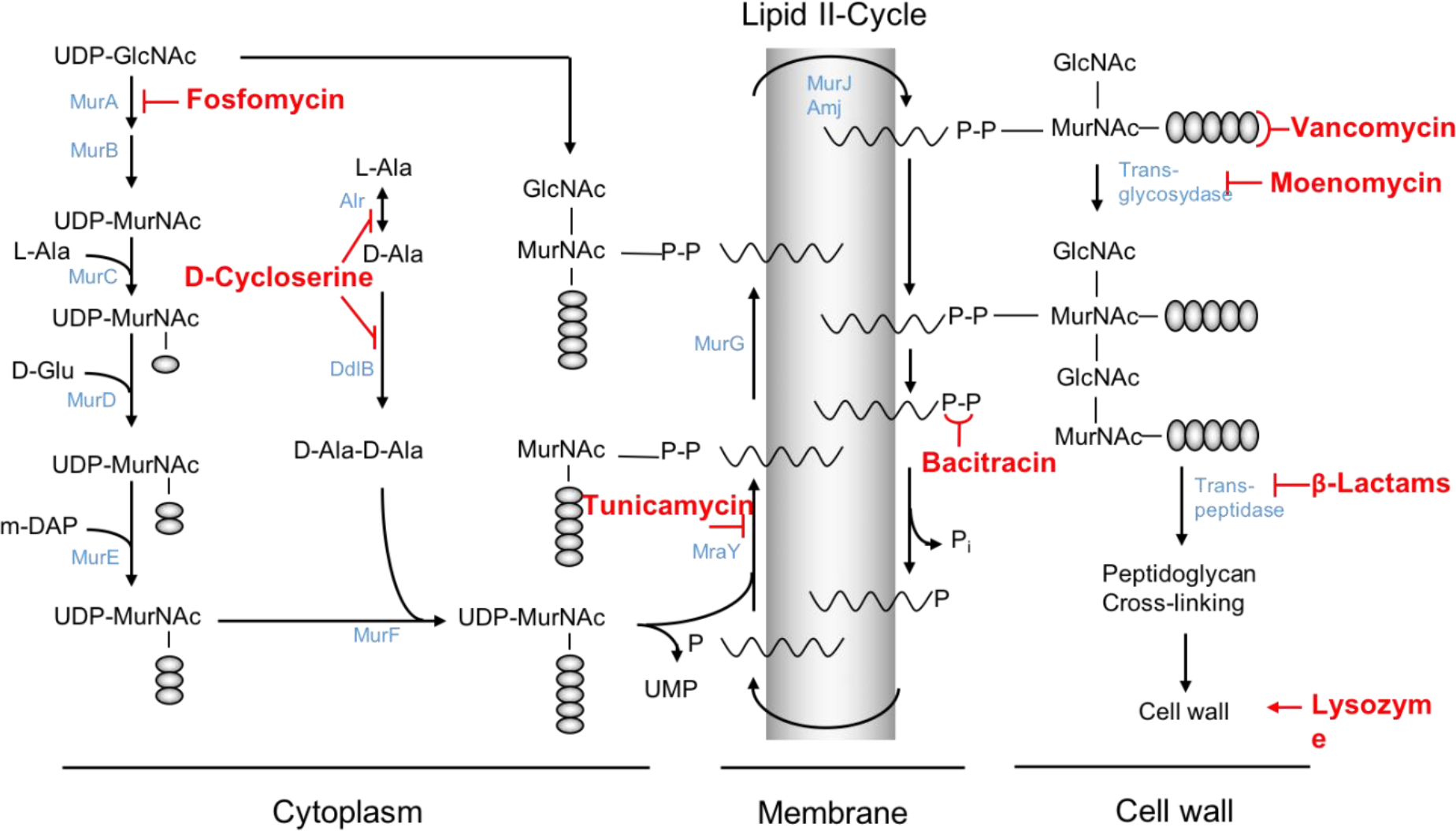
The peptidoglycan biosynthetic pathway showing sites of action of the compounds applied in the study. The pathway involves three stages as described in the main text. Cell envelope-active compounds applied in this study are given and placed next to the steps they inhibit. Some of the enzymes involved in the cell wall biosynthesis are shown in light blue next to the corresponding catalytic steps. Lipid II is the GlcNAc-MurNAc- pentapeptide covalently linked to the lipid carrier undecaprenol (symbolized by the waved lines) via pyrophosphate ester bridge. Abbreviation: GlcNAc, N-acetyl-glucosamine; MurNAc, N-acetyl-muramic acid. Amino acids are symbolized by small grey circles. This figure was generated based on (2, 3) with modifications.

Lastly, lysozyme is an enzyme that kills bacteria by hydrolyzing the 1,4-β-link between MurNAc and GlcNAc residues in the PG [22].

### Protecting the wall: Cell envelope stress response (CESR) in *Bacillus subtilis*

Because of its vital role and numerous potential threats, the integrity of the cell envelope is closely monitored. Countermeasures, e.g. against antibiotic action, can then be implemented to cope with stress before irreversible CW damage can occur. Collectively, these measures are termed the cell envelope stress response (CESR). In the past 25 years, the underlying regulatory network has been extensively studied in the Gram-positive model organism *B. subtilis* (summarized in [23, 24]). At least eight regulatory systems from two major signaling principles, extracytoplasmic function sigma factors (ECFs) and two-component signal transduction systems (TCSs), are involved in mediating the CESR in this organism. Four of the seven ECFs have been directly linked to counteracting stress or damage caused by CW antibiotics, of which σ^W^ and σ^M^ play a predominant role in providing a more general resistance against cell envelope damage, while σ^X^ and σ^V^ are specific for membrane perturbations or lysozyme challenge, respectively [25]. σ^W^ controls a large ‘antibiosis’ regulon, with a significant number of its 60-90 target genes encoding functions implicated in antibiotic resistance. Accordingly, a *sigW* mutant is more sensitive to fosfomycin, pore-forming lantibiotics (such as nisin) and a number of antimicrobial peptides produced by *Bacillus* sp. Taken together, σ^W^ is induced by envelope stress and protects the cell against antibiotics and bacteriocins, especially if they have membrane-disruptive properties, e.g. by altering membrane lipid composition [26]. σ^X^ contributes to the resistance against cationic antimicrobial peptides (AMPs) by altering cell surface properties [27]. Accordingly, a *sigX* mutant is more sensitive to cationic AMPs such as nisin, which were not included in this study. The *dltA* operon (D-alanylation of teichoic acid) and *pssA* operon (biosynthesis of phospholipid) were previously shown to be most strongly activated by σ^X^ [27]. Both of the systems decrease the net negative charge of the cell envelope, reducing AMPs binding. σ^V^ is strongly and specifically induced by lysozyme and its induction provides lysozyme resistance [28]. Despite some regulatory overlap with σ^X^ at the level of promoter recognition, σ^V^ stress response presumably has evolved to defend against lytic enzymes.

In contrast to the CESR functions described for the three ECFs above, σ^M^ plays a much more fundamental role in modulating the core PG biosynthesis and cell division machinery of *B. subtilis*, thereby maintaining the integrity of the CW in the presence of CES. While a *sigM* mutant is highly sensitive to β-lactams, its anti-σ factor is encoded by an essential gene, indicating that the cell cannot tolerate a dysregulation of essential processes caused by the resulting upregulation of the σ^M^ regulon [25].

Four out of 32 TCSs are directly involved in mediating the CESR of *B. subtilis:* LiaSR, BceRS, PsdRS, and ApeRS [29]. The TCS LiaRS of *B. subtilis* was originally named for lipid II cycle interfering <u>a</u>ntibiotic response regulator and <u>s</u>ensor. Accordingly, the LiaR target operon, *liaIH*, can be strongly induced in the presence of the antibiotics that interfere with the lipid II-cycle [30]. In addition, membrane-active compounds such as daptomycin also activate the Lia response [31, 32], most likely by indirectly interfering with lipid II. But despite extensive studies, the true nature of the signal provoking the Lia system and also its biological role remain poorly understood. BceRS, PsdRS and ApeRS are three paralogous TCSs that are specifically induced by and mediate resistance against CESR. They are functionally and genetically associated with ABC transporters, and together form a unique type of AMPs detoxification modules that are widely conserved in *Firmicutes* bacteria [33]. AMPs bind to and are sensed through the cognate ABC transporters, which indirectly activates the TCS. In response, the corresponding ABC transporter genes are strongly induced and their gene products remove the AMP from the cell surface, thereby mediating resistance [34, 35]. All of these systems show a high substrate-specificity [36].

Two additional TCSs have regularly been associated with the CESR of *B. subtilis*. The TCS CssRS (<u>c</u>ontrol <u>s</u>ecretion <u>s</u>tress regulator and <u>s</u>ensor) controls the cellular responses to protein secretion stress in *B. subtilis* [37]. The stress of high-level production of secretory proteins mounts the CssRS-dependent induction of *htrA* and *htrB*, which encode extracellular membrane-anchored quality control proteases [37, 38]. The TCS WalRK orchestrates CW homeostasis in *B. subtilis* and is essential for its viability [39]. It was originally characterized in *B. subtilis*, but is widely conserved in, and specific to *Firmicutes* bacteria, including a number of important pathogens [39–43]. In *B. subtilis*, WalRK controls a set of genes that are either activated or repressed by the WalR response regulator [39, 44, 45]. When CW metabolism is particularly active, e.g. during the exponential growth phase when cells are rapidly growing and dividing, the WalRK system is highly activated. As a result, genes positively regulated by WalR, such as *cwlO* and *lytE* (encoding the co-essential D,L-endopeptidase type autolysins LytE and CwlO involved in PG elongation), *yocH* (peptidoglycan amidase) and *ftsAZ* (cell division), show a higher expression level to ensure high CW plasticity for cell growth [46–48]. In contrast, genes negatively regulated by WalR, such as *iseA* (inhibitor of LytE and CwlO) and *pdaC* (peptidoglycan deacetylase C), are repressed [39, 49, 50]. In non-diving cells (stationary phase), WalRK activity is tuned down. Repressed genes of the WalR regulon will be released from repression, while the activated genes are transcriptionally downregulated. As a consequence, CW turn-over is reduced, in line with the arrested CW growth and halted cell division.

### Profiling the CESR of *B. subtilis*

While numerous studies have been performed in the past to analyze the transcriptional response of *B. subtilis* to individual CW antibiotics (summarized in [51]), many are from the early days of transcriptomics and are often of low quality due to the experimental procedures, parameters and platforms applied. The sensitivity and dynamic range of early macro- and microarrays were far from what can be resolved with current approaches. But even more importantly is the choice of conditions for stress response experiments. Sublethal antibiotic concentrations and short incubation times between induction and harvest are the two most critical parameters to ensure that only the specific, that is the primary, transcriptional CESR is monitored [51]. Signal transduction and gene regulation are inherently fast processes and full responses to antibiotic challenge can be monitored already after 3-5 min [52, 53]. In contrast, higher antibiotic concentrations (at or even above the MIC) and prolonged exposure to the antibiotic (30-60 mins were often applied) leads to an accumulation of cellular damage and increasingly unspecific transcriptomic signatures. In the worst case, the specific primary responses are masked or already shut off [51]. Variations in experimental procedures also hamper a meaningful comparison between different transcriptome profiles, thereby ultimately preventing to gather a comprehensive picture of the CESR response, when challenged with different CW antibiotics. Moreover, virtually all previous studies refer to the initial *B. subtilis* genome sequence [54], which contained numerous errors and missed many genomic features, such as small non-coding RNAs that were only uncovered much later in the course of updating the genome sequence [55].

This study aims at revealing the genome-wide transcriptional response of *B. subtilis* to a set of PG synthesis inhibitors and providing a comprehensive picture of the CESR of *B. subtilis* from a transcriptomic point of view. A set of compounds that interfere with all three stages of PG biosynthesis, the initial intracellular steps (fosfomycin and D-cycloserine), the membrane-associated lipid II-cycle (tunicamycin, bacitracin, vancomycin) as well the final extracellular steps (moenomycin and penicillin G) (Fig. 1) were used at sublethal concentrations. Lysozyme was also included as an agent that destroys the already made murein sacculus. In order to analyze the performance of our profiling efforts and validate the data, we applied in parallel two independent current methods of transcriptional profiling: RNAseq and the latest generation of *B. subtilis* tiling arrays, which had previously been established as the gold standard for studying gene expression levels on a global scale [56]. Each stimulon was carefully dissected and comparatively analyzed to uncover the role of ECFs and TCSs in the CESR. ECF regulons were refined and a set of *B. subtilis* whole-cell biosensors were constructed and evaluated for their functionality.

## Results

### Experimental design, data processing and identification of CESR-induced genes

Initially, the inhibitory activity of the eight antimicrobial compounds was carefully analyzed on wild type *B. subtilis* cells growing in LB medium at 37°C, in order to determine the appropriate sub-lethal concentrations to be used for our transcriptomic experiments (Fig. S1). Cells at mid-exponential growth phase (OD600 ≈ 0.4) were then treated for 10 min, and samples were collected for RNA extraction, cDNA library preparation and either RNA sequencing or tiling array hybridization. The resulting raw sequencing reads and hybridization patterns, respectively, were analyzed to identify compound-specific and common changes of gene expression caused by the eight antibiotics, using untreated samples as negative control (see Experimental Procedure for details).

The mapping of expression signals was referred to the annotation file “BSGatlas_v1.0.gff” from BSGatlas (https://rth.dk/resources/bsgatlas/), which contains 4773 generic features including coding- and non-coding genes, UTRs, transcripts, TSSs, and terminator structures [57]. SubtiWiki (http://subtiwiki.uni-goettingen.de/) was used as the “official” reference for both gene annotations and the definition of regulons, as this platform is manually curated to continuously incorporate and update the latest findings on *B. subtilis* [58].

We first applied a threshold of four-fold change of gene expression in at least one treatment condition as an initial filter. This resulted in 307 genes, 66 ncRNAs, 14 new RNA features and 22 novel transcripts that were differentially expressed in the RNAseq dataset as compared to a non-treated control (Table S1) and 212 genes and 84 ncRNAs differentially expressed in the tilling array dataset (Table S2). This corresponds to 8.6% and 4.4% of all expressed genes in our RNAseq and tilling arrays analysis, respectively. Next, the transcripts with very low basal expression (less than 10 transcriptional reads on average for RNAseq (Table S1) or a level of expression under 9 for tiling arrays) were manually removed to avoid irrelevant fold-change of gene expression. In addition, the 20 rRNAs were also removed because of their high and consistent expression levels (Fig. S2). The remaining 327 differentially expressed transcripts that include 298 genes, 16 ncRNAs, 7 new RNA features and 6 novel transcripts were then subjected to in-depth analyses. Hierarchical clustering analysis was performed using the Heatmap2 package in R program. This unsupervised clustering algorithm divided the large list of differentially expressed genes into 11 clusters of similar patterns (C1 to C11, Table S3). These clusters correlate with (combinations of) distinct regulons and allowed visualizing the specific expression patterns within each stimulon, thereby enabling the analysis of specific expression signatures based on the activation of distinct signaling pathways (Fig. 2, RNAseq and Fig. S3 tilling array). Graphical representations of the individual stimulons are provided in Fig. 3. For reasons of clarity and simplicity, only RNAseq data are shown hereafter in the main figures, while the corresponding tiling array data are provided in supplemental material (tables S5,7,9,11,13,15,17,19). The good correlation of both approaches is illustrated in the regulon-specific expression data provided in Fig. 4-Fig. 6.

**Fig. 2.**
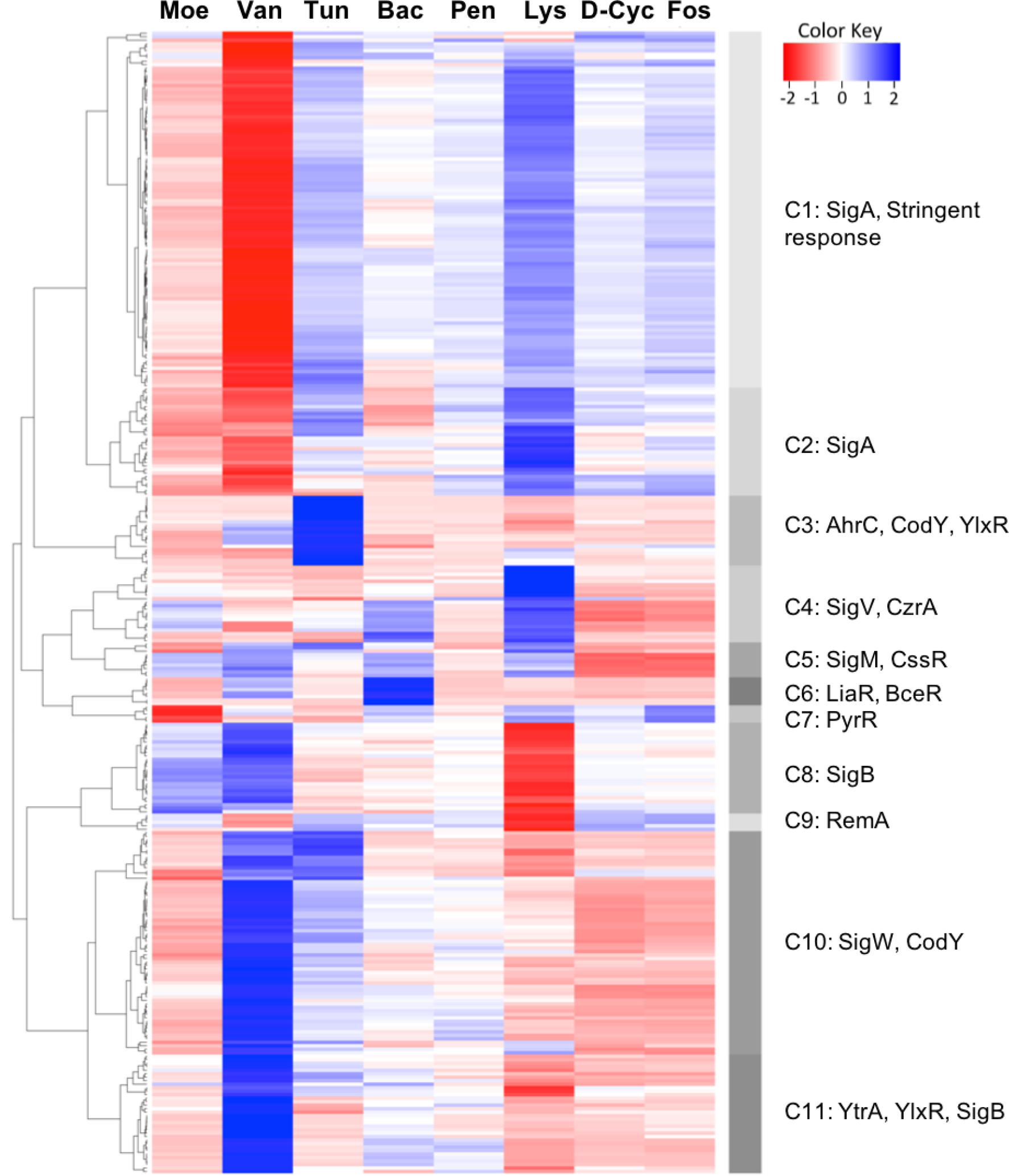
Hierarchical clustering analysis of the transcriptional profiles of *B. subtilis* in response to cell envelope- active compounds (RNAseq).The heatmap was generated using “Heatmap.2” package in R program based on the log2 fold-change of the 327 differentially expressed genes. Side bar marks the 11 Clusters generated during clustering analysis. The C in front of the numeral denotes “Cluster”, with the major regulons in that cluster displayed. Each column represents the corresponding stimulon as indicated on the top in three letters. Each row represents the expression of one gene across the stimulons. The Color Key square on the top left corner indicates that the highest gene expression level is colored in blue, while the lowest gene expression level is in red.

**Fig. 3.**
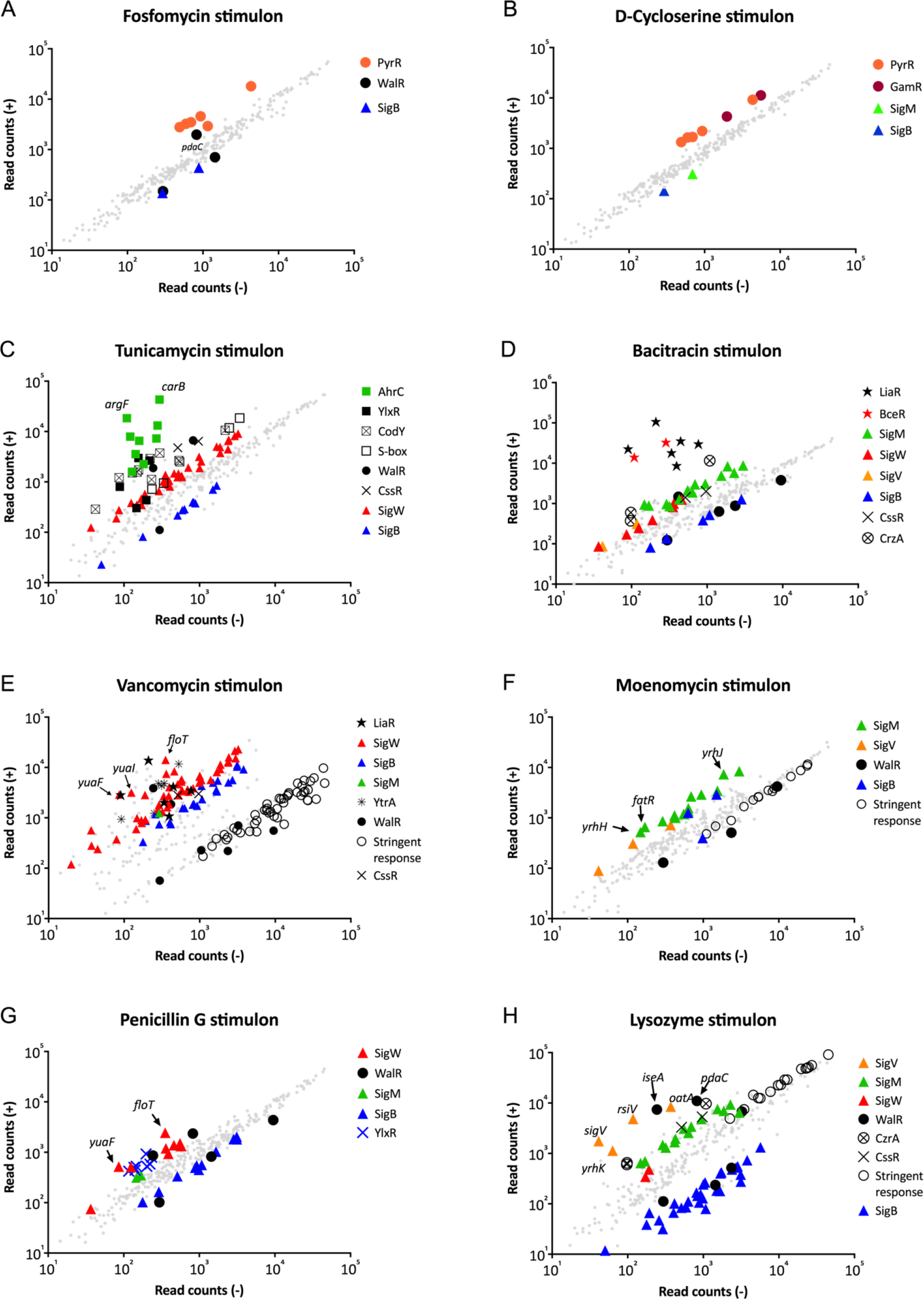
Graphic presentation of the stimulons. Each graph represents the read counts of the genes in the treated sample (y-axis) against that in the untreated sample (wild type, x-axis). Most of the genes induced or repressed by at least 2-fold are highlighted using different symbols as indicated on the graph for each stimulon, with the corresponding regulators shown on the right side. Other genes are represented as smaller grey solid cycles. The read counts of each gene represent the mean value from the triplicates of each treatment.

**Fig. 4.**
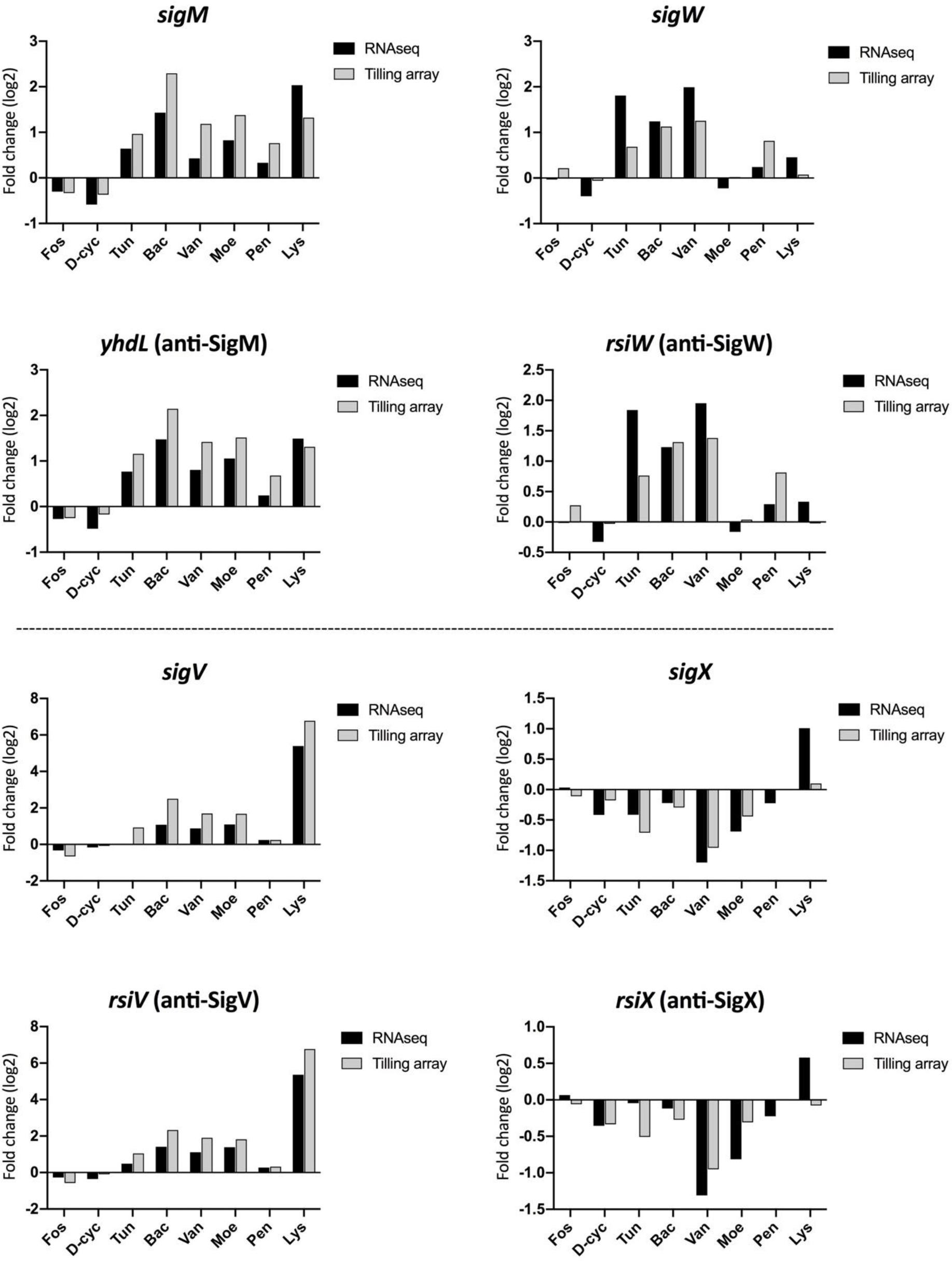
Induction profiles of *sigW*, *sigM*, *sigV* and *sigX* by the CESs.The fold-change of each ECF σ factor represents the mean value from the triplicates of each treatment in RNAseq.

### Inhibition of membrane-anchored steps of PG synthesis induces a strong CESR, in contrast to cytoplasmic and extracellular steps

In both the RNAseq and tiling array data, CW antibiotics inhibiting the early cytoplasmic steps (fosfomycin and D-cycloserine) and the extracellular crosslinking reactions (moenomycin and penicillin G) did not trigger a pronounced transcriptional response. In contrast, antibiotics interfering with the membrane-anchored steps (bacitracin, tunicamycin and in particular vancomycin) provoked strong and specific responses (Fig. 2, Fig. 3 and **Erreur ! Source du renvoi introuvable.**3). This dominant feature may reflect the membrane-proximate perception of envelope stress by the TCSs and ECFs involved (see below). Lysozyme, which actively damages the CW, was also a strong CESR inducer, with a transcriptional signature that seemed to almost anti-correlate with the vancomycin stimulon. The top five genes triggered by each stimulus are present in Table 1. All genes showing a fold-change difference of at least two (fosfomycin, D-cycloserine, moenomycin and penicillin G) or five (tunicamycin, bacitracin, vancomycin and lysozyme) have been summarized in the compound-specific 4 to Table S19. Each of the eight antibiotics caused between 0 and 275 genes or ncRNAs to be differentially expressed as compared to non-treated control cells. 12/1 genes were differentially expressed in response to fosfomycin, 11/0 to D-cycloserine, 103/18 to tunicamycin, 81/61 to bacitracin, 275/110 to vancomycin, 79/0 to moenomycin and 131/22 to lysozyme in the RNAseq/tilling array data.

### The stimulons: antibiotic-specific transcriptional profiles of *B. subtilis*

**Fosfomycin**, which targets the first committed intracellular step of PG precursor biosynthesis, the conversion of UDP-GlcNAc to UDP-MurNAc by MurA (Fig. 1), provoked only a minor response in *B. subtilis* (Fig. 3A). Only the *pyr* operon (Fig. 2, cluster 7), involved in pyrimidine metabolism, was induced above 5-fold (Fig. 3A, and Table S4 and Table S5). A weak induction of *pdaC* and *iseA*, negatively controlled by WalR (Fig. 6), as well as genes associated with glucosamine utilization (*gamAP*) are in line with responding to an inhibition of PG precursor biosynthesis. PdaC is a PG deacetylase that confers lysozyme resistance [49], while IseA acts as an inhibitor of PG hydrolases that reduces the rate of antibiotic-induced cell death [50]. No ECF-dependent gene expression was observed in response to fosfomycin, nor did this compound trigger any of the four typical TCS involved in the CESR of *B. subtilis*. **D-cycloserine** inhibits the formation of the dipeptide D-alanyl-D-alanine (D-Ala-D-Ala) (Fig. 1). The response of *B. subtilis* to D-cycloserine was even weaker, but otherwise comparable to that to fosfomycin. The *pyr* and *gamAP* operons were weakly induced (Fig. 3B and Table S6). Like fosfomycin, D-cycloserine is known as a σ^W^ inducer [52] but this activation was not detected in our experimental conditions, after 10 min of treatment.

**Tunicamycin** targets the first membrane-associated step of PG biosynthesis by preventing the formation of lipid I from UDP-MurNAc-pentapeptide [59]. Additionally, tunicamycin also interferes with the formation of wall teichoic acids [14]. Previously, tunicamycin was shown to weakly induce σ^ECF^ and the Lia system [30, 52, 60, 61], which could be confirmed by our transcriptomic study. ECF-dependent gene expression was primarily orchestrated by σ^W^ (Fig. 3C), but – to a weaker extent – also by σ^M^ (Fig. 4). A moderate activation of the Lia system by tunicamycin was also detected (Fig. 5), which is consistent with an earlier study [30]. Surprisingly, many AhrC- and CodY-controlled operons related to amino acids metabolism (e.g. biosynthesis of arginine, leucine, branched-chain amino acids, methionine, and cysteine) were amongst the most highly induced genes (Fig. 2 cluster 3, Fig. 3C and Table S8 and Table S9). Finally, the WalR-dependent genes *pdaC* and *iseA*, also induced by fosfomycin and D-cycloserine, and the CssR-dependent secretion stress-inducible genes *htrA* and *htrB* were also activated (Fig. 3C, Fig. 5, Fig. 6, Table S8 and Table S9). Tunicamycin therefore strongly triggered amino acid metabolism genes and, to a lesser degree, the induction of σ^W^ and σ^M^ regulons, as well as affecting the TCSs LiaRS, WalRK, and CssRS.

**Fig. 5.**
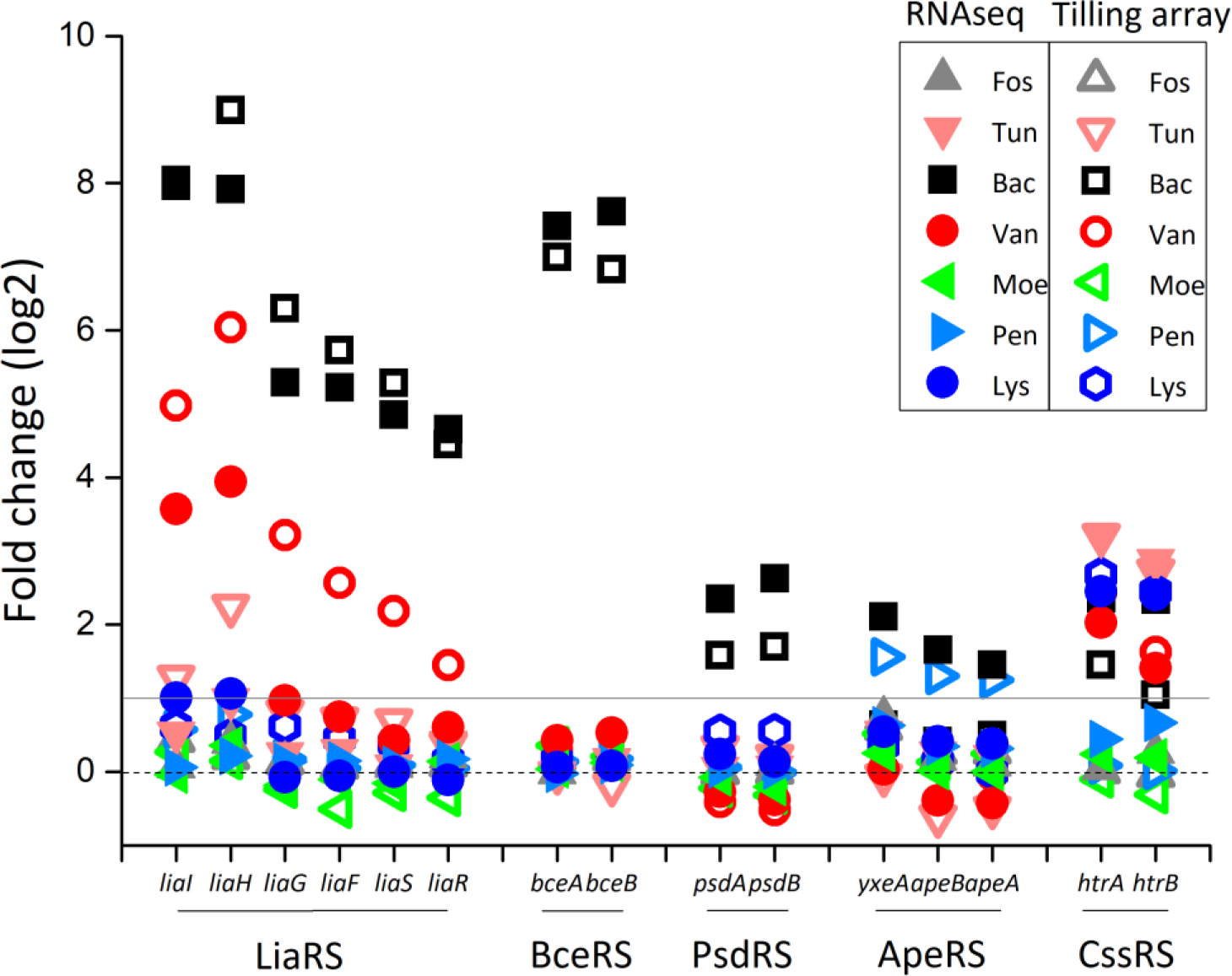
Transcriptional response of two-component systems under cell envelope stresses. The operons regulated by the TCSs are indicated on the x-axis, with the corresponding TCS shown below. Each stack shows the DGE of each gene under different treatments, with the special conditions highlighted by distinct colors or symbols.

**Fig. 6.**
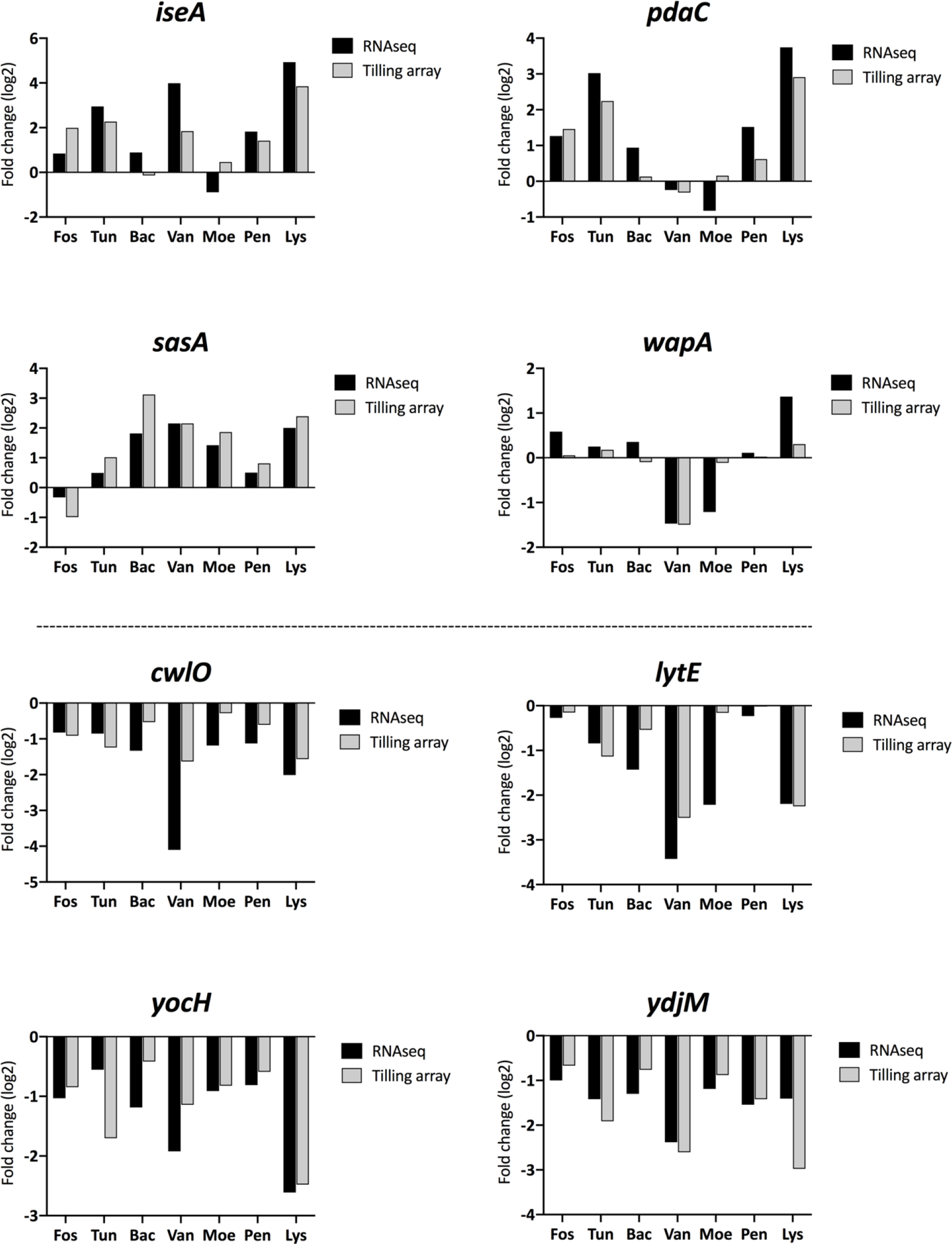
Transcriptional response of WalR regulon to cell envelope-active compounds. The four graphs on the top half represent the induction profiles of the four genes repressed by WalR, and the four graphs below the dashed line represent the induction profiles of four of the genes activated by WalR.

**Bacitracin** prevents the recycling of the lipid carrier undecaprenol (Fig. 1). In line with previous studies [53], bacitracin strongly activated the two operons *liaIH*-*liaGFSR* and *bce*AB, which are under control of the TCS LiaRS and BceRS, respectively (Fig. 2 cluster 6, Fig. 3D, Table S10 and Table S11). The *bce*AB operon encodes the ABC transporter BceAB, which acts as the primary bacitracin resistance determinant, while LiaIH provide a secondary layer of resistance [24, 53, 62] (Fig. 5). As observed previously, two BceRS-paralogs, PsdRS and ApeRS, were also weakly induced, presumably through cross-activation by BceRS [52, 62]. The ECF-dependent response to bacitracin was less pronounced than the TCS-mediated response and primarily mediated by σ^M^ and to a lesser extent by σ^W^ and σ^V^ (Fig. 4), in line with previous observations [53]. Together, these ECFs control induction of *bcrC*, encoding a second undecaprenyl pyrophosphate phosphatase that functions as a bacitracin resistance determinant [60, 63].

Since bacitracin is complexed with Zn(II) ions, which are also required for this antibiotic to be biologically active [64], bacitracin treatment also activated the CrzA-mediated toxic metal ion stress response by inducing *cadA* and the *czcD-czcO* operon (Table S10 and Table S11), which mediate resistance against them [65]. All of the above responses have been observed and characterized previously [53]. In contrast, induction of *yrhH*-*fatR*-*yrhJ* (fatty acid biosynthesis) and *hisZGDBHAFI* (histidine biosynthesis) was observed for the first time in this study (Fig. 8, Table S10 and Table S11). **Vancomycin** inhibits PG biosynthesis by binding to the D-alanyl-D-alanine dipeptide terminus of externalized Lipid II, thereby blocking glycan polymerization and cross-linking [18]. Vancomycin was already known as a strong inducer of the CESR in *B. subtilis*, in particular the σ^W^ regulon [52]. Our results (Fig. 2 cluster 10, Fig. 3E and Fig. 7) are consistent with these findings. The LiaRS-regulated *liaIH- liaGFSR* operon and the σ^W^ regulon predominated the primary response of *B. subtilis* to vancomycin (Fig. 3E, Table S12 and Table S13). σ^M^ and σ^V^ were also activated, but to a lesser degree (Fig. 4).

**Fig. 7.**
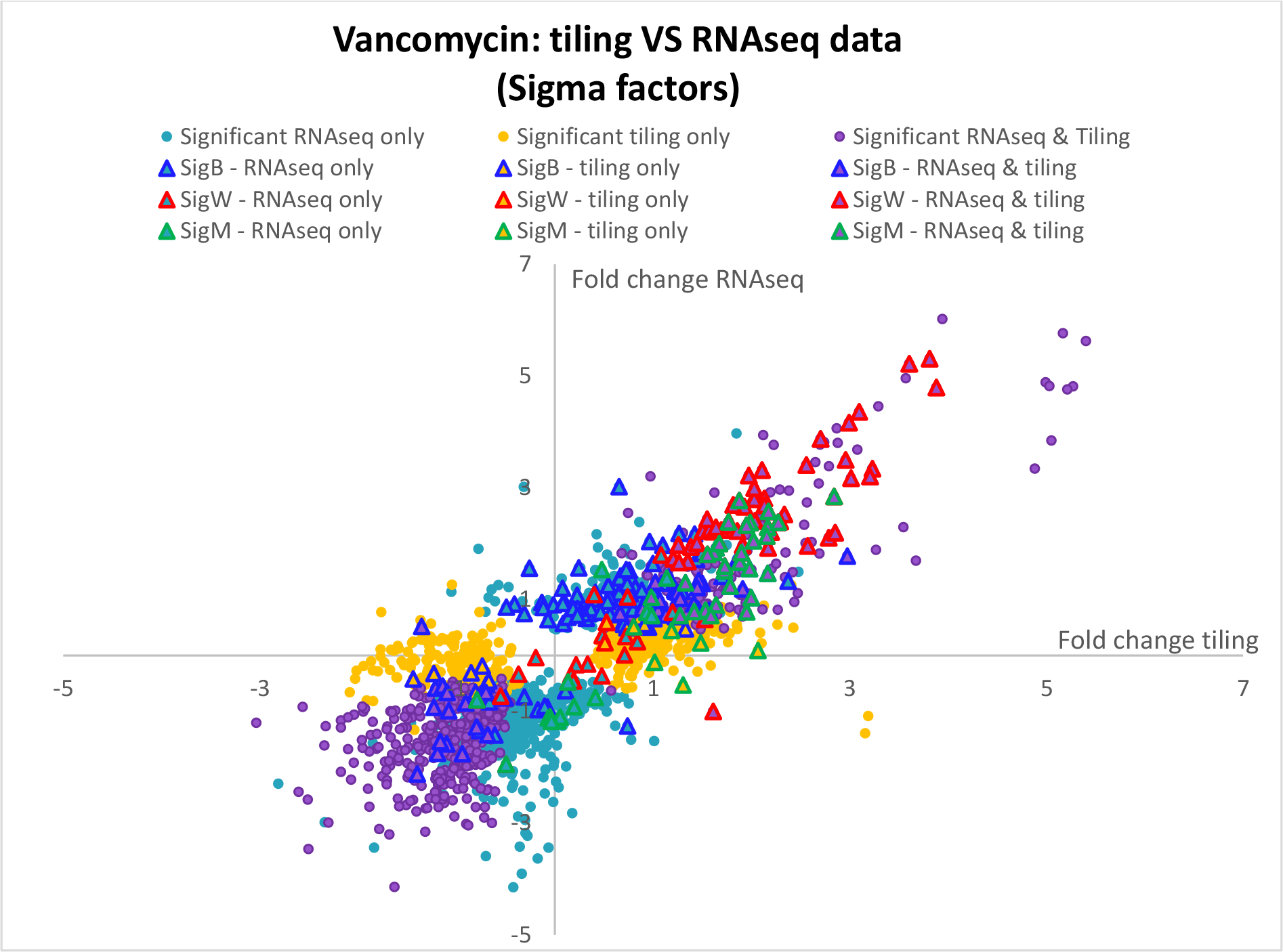
Vancomycin stimulon (RNAseq vs Tilling array)

**Fig. 8.**
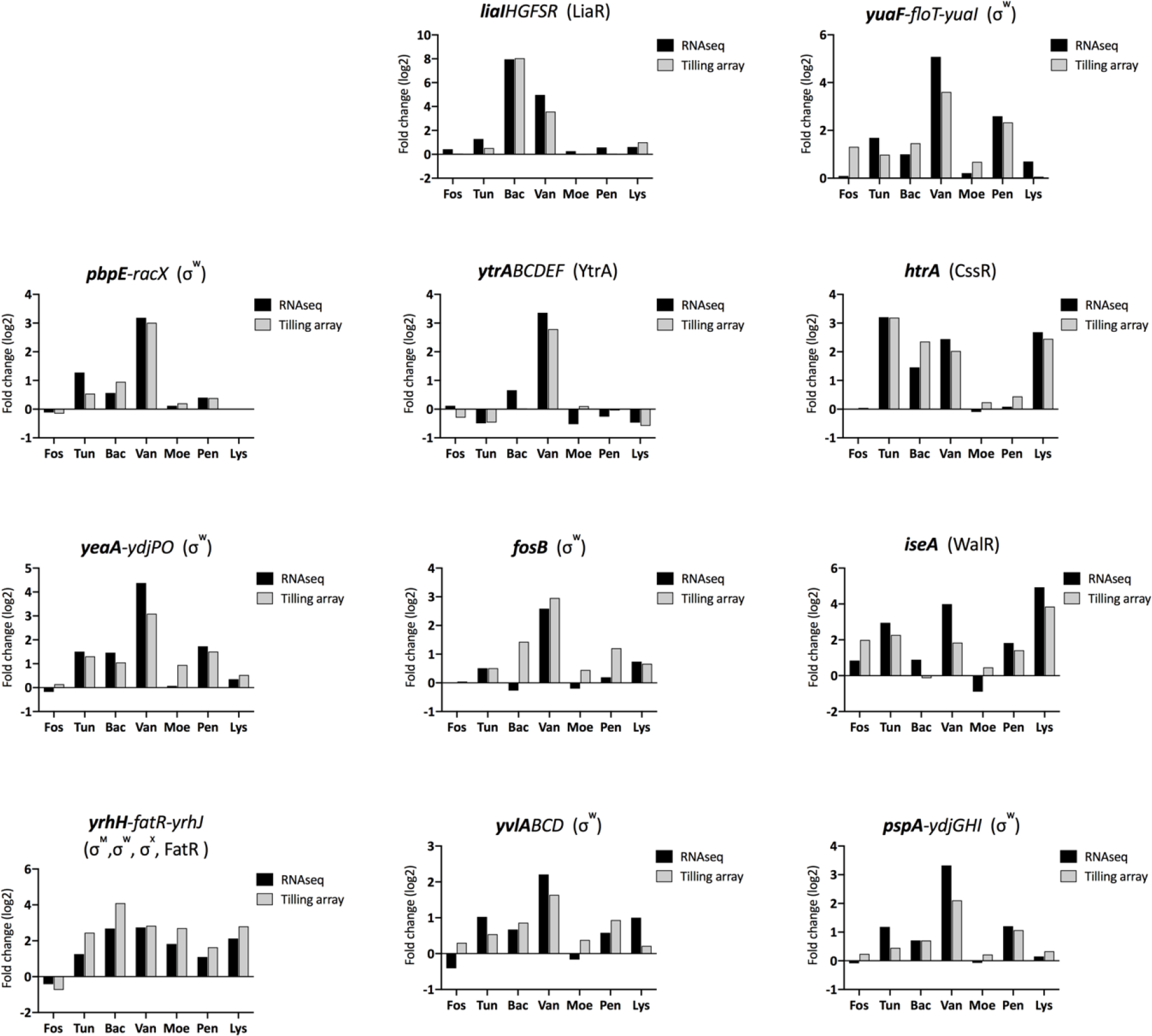
Expression patterns of CESR marker genes. In case of operons, the figure represents the induction profile of the gene of the operon marked in bold (usually the first gene of the operon). The corresponding regulator(s) for each gene or operon are displayed in parentheses.

Several signature genes of the vancomycin stimulon are related to inhibition of both PG synthesis and hydrolysis. The WalR-controlled PG hydrolases were inhibited by induction of the *iseA* gene (modulating autolysins activity) and repression of the co-essential *lytE* and *cwlO* (encoding DL-endopeptidases), and of *ydjM* (encoding a CW-associated protein). In parallel, genes involved in PG synthesis (e.g. *murE-mraY-murD-spoVE operon*, *mbl, dacA*) were also repressed. Interestingly, *yrhH*- *fatR*-*yrhJ* was strongly induced, as observed for bacitracin. The CssRS-dependent protein quality control genes *htrAB* were also induced by vancomycin, as by many other CW antibiotics (Fig. 5).

In contrast to the other antibiotics analyzed in this study, different vancomycin concentrations were used between RNAseq (1 µg/ml) and tiling array (0.25 µg/ml) studies. While this 4-fold difference in antibiotic concentration did not change the overall picture of the primary CESR (the same regulons were identified in both vancomycin stimulons), quite a number of its genes showed differences in the overall induction strength, which was mostly higher at 1 µg/ml (Fig. 4 and Fig. 8). More importantly, it resulted in significant differences in the secondary, less specific, global responses (Fig. 7, Table S12 and Table S13): The σ^B^-dependent general stress response was weakly activated at 1 µg/ml vancomycin in the RNAseq experiments, indicating that the primary and more specific responses did no longer provide enough protection at this higher antibiotic concentration [66]. Moreover, the stringent response was more severely affected at higher vancomycin concentrations, as indicated by the repression of many ribosomal protein-encoding genes in the RNAseq data (Fig. 3E and Table S12).

Taken together, vancomycin triggers a very strong CESR in *B. subtilis*, which is largely mediated by the TCS LiaRS and WalKR, σ^W^ and (at higher vancomycin concentrations as observed in RNAseq) the σ^B^- dependent general stress response. The results obtained so far demonstrate that global expression studies on antibiotic stress are rather robust to different technological platforms, but strongly affected by the antibiotic concentrations applied. Secondary, global responses are usually induced at higher concentrations by the accumulating damage caused by antibiotic action [51].

**Moenomycin** targets the glycosyltransferase activity of aPBPs [20]. Similar to the inhibition of the intracellular steps of PG biosynthesis, only a weak transcriptional response is observed for this antibiotic. The *pyr* operon (involved in uracil metabolism) was the only locus to be repressed ≥ 5-fold, while no gene was induced ≥ 5-fold, in line with a previous study [67] (Fig. 3F, Table S14 and Table S15). A weak activation of σ^M^ and σ^V^ was nevertheless detectable (Fig. 3 and Fig. 4). Interestingly, the strongest induction was observed for the *yrhH-fatR-yrhJ* locus, which is jointly regulated by σ^M^, σ^W^, σ^X^ and FatR. Other strongly induced genes were also co-regulated by several ECFs (Table S14 and Table S15). Of the different ECF σ factors, σ^M^ was reported to be the only one contributing to moenomycin resistance in *B. subtilis*: A *sigM* deletion strain was much more sensitive to moenomycin than any other σ^ECF^ mutation and only the overexpression of σ^M^ in the Δ*7ECF* mutant was able to restore the resistance of *B. subtilis* to this compound [68]. But overall, inhibiting the TG activity of aPBPs by moenomycin only triggered a minor response in *B. subtilis*.

**Penicillin G** and related β-lactams inhibit the TP activity of so-called penicillin-binding-proteins (PBPs). Again, the response of *B. subtilis* to this extracellular inhibitor of PG biosynthesis was not very pronounced (Fig. 2, Fig. 3G, Table S16 and Table S17). The σ^W^ regulon was activated, with the *yuaF*- *floT*-*yuaI* operon being the only one induced ≥ 5-fold. σ^M^ was also slightly induced by penicillin G, while σ^X^ and σ^V^ were not responsive (Fig. 3G and Fig. 4). The *apeAB*-*yxeA* operon, which is controlled by the TCS ApeRS, was surprisingly induced 2.5 fold (Fig. 3G and Fig. 4) since ApeRS usually responds to antimicrobial peptides of eukaryotic origin. Nevertheless, our data indicate that penicillin G is a weak inducer of the CESR of *B. subtilis*, in line with previous reports [69].

### Lysozyme stimulon

Lysozyme kills bacteria by cleaving the β-1,4-glycosidic bonds between the MurNAc and GlcNAc, resulting in cell lysis [70]. It is a strong inducer of σ^V^ through direct binding of lysozyme to the membrane-anchored anti-σ factor RsiV [28, 71]. The activation of σ^V^ confers lysozyme resistance through OatA-dependent PG O-acetylation (encoded in *sigV*-*rsiV*-*oatA*-*yrhK* operon) and DltABCDE-dependent D-alanylation of teichoic acids [71, 72], which is controlled by σ^X^, σ^V^ and σ^M^. Indeed, we observed a >40-fold induction of the *sigV* operon upon lysozyme addition (Fig. 3H and Table 1), while the *dlt* operon was only induced approximately two-fold (Table S18 and Table S19). Additional ECFs were also induced by lysozyme, including σ^M^, σ^X^ and σ^W^ (Fig. 3H and Fig. 4). A strong induction was again observed for *iseA* and *pdaC* (approx. 30- and 13-fold, respectively). Interestingly, the CzrA- controlled *czcD*-*czcO* operon and *cadA* gene, which normally respond to metal ion stress [65], were strongly activated by lysozyme, as already observed for bacitracin. While the induction by bacitracin was due to the Zn(II) ions coordinated by this antibiotic, the reason for the CzrA response to lysozyme will require further investigations. Other genes induced strongly and exclusively by lysozyme include the *maeA*-*ywkB* operon and *maeN* gene, which are controlled by TCS MalRK involved in malate utilization [73], and the CssR-controlled genes *htrA* and *htrB*, which were also induced by tunicamycin. Despite the severe CW damage caused by lysozyme, the σ^B^-dependent general stress genes was strongly repressed by lysozyme, while stringent response-associated genes were upregulated. Taken together, our data show that disruption of CW by lysozyme triggered a strong and complex response with the significant activation of σ^V^ and σ^M^, as well as the involvement of TCS WalRK.

### Refinement of the ECF σ factor regulons

The σ^ECF^-dependent antimicrobial resistance network constitutes one of the two major regulatory routes mediating the CESR in *B. subtilis*. Of the seven σ^ECF^ factors encoded in the genome of *B. subtilis*, four (σ^W^, σ^M^, σ^X^ and σ^V^) are well-understood in terms of their roles in cell envelope homeostasis and antibiotic resistance. Their regulons have been extensively investigated and are well determined [25, 74, 75]. Remarkably, a significant extent of regulatory overlap was observed due to the similarities of promoter sequences recognized by those σ^ECF^ factors [60, 76–78]. The resulting functional redundancy [79] still poses a challenge in determining the contribution of individual σ^ECF^ factors to the expression of genes assigned to multiple regulators. We therefore attempted to refine the σ^W^, σ^M^, σ^X^ and σ^V^ regulons by integrating the comprehensive transcriptional profiles generated in this study with the previously established detailed information on each regulon and their regulatory overlap. Towards this goal, we (i) determined the major regulator of genes under control of multiple σ^ECF^ factors based on the distinct expression pattern of each regulon (Fig. 4), (ii) reevaluated the members of the four σ^ECF^ regulons according to their response to the different stresses (Table 2), and (iii) searched for novel candidates for the four σ^ECF^ regulons via hierarchical clustering analysis (Fig. S45 and Table S2020). Based on this combined analysis, five different groups of ECF-dependent genes could be identified (Table 2).

Group I includes genes that appear to have σ^M^ as their major regulator. They were induced by lysozyme, bacitracin and moenomycin. Some genes, such as *bcrC* and *divIC*, are regulated by additional ECFs.

Group II genes are primarily controlled by σ^X^. This group of genes was only activated by lysozyme and repressed by vancomycin and sometimes also other CW antibiotics, such as bacitracin or moenomycin (Table 2). The overall response of σ^X^-dependent genes appeared to be very weak to the stresses applied in this study.

Group III genes are exclusively regulated by σ^V^. Most of the σ^V^-dependent genes are also controlled by σ^X^, σ^M^, and σ^W^ [77], with only the *sigV* operon being exclusively regulated by σ^V^. In this study, the *sigV* operon was most strongly induced by lysozyme, but also weakly by bacitracin, vancomycin and moenomycin (Table 2).

Group IV genes are controlled by σ^M^, but may also be significantly regulated by other ECFs. Genes from within this group showed a broader inducer spectrum. While they are also partially controlled by σ^M^ but are more strongly induced by vancomycin than the preferentially σ^M^-dependent genes of Group I. Notably, they all have additional regulator(s) other than ECFs, such as σ^B^, FatR, Spx, and WalR (Table 2). σ^W^ seems to play only a secondary role in regulating genes of group IV. Genes in this group perfectly match the previously complied list of genes that were σ^W^-dependent but not significantly down- regulated in a σ^W^ mutant [80].

Finally, Group V includes all genes primarily (mostly exclusively) regulated by σ^W^ (Table 2), with the exception of *yceCDEFGH* operon, which is also regulated by σ^M^, σ^X^ and σ^B^ [74, 75, 81, 82]. The sequence alignment revealed that the promoters of these genes share characteristic sequence motifs at the -35 and -10 regions (Fig. S5). In particular, the -10 region with “CGTA” motif is clearly distinct from the corresponding regions of other ECF-target promoters in *B. subtilis*, which more frequently show a “CGTC” motif.

Taken together, our analysis provides a comprehensive picture of ECF-dependent gene expression. While it confirmed most of the known ECF-target genes, it also identified potentially novel members of the σ^W^ regulon. The comprehensive analysis of the distinct expression patterns of the ECF regulons under different CESs enabled us to distinguish the partially overlapping ECF regulons and allowed determining the major regulators for the genes co-regulated by different ECFs.

### The role of TCSs in mediating CESR of *B. subtilis*

Four TCSs represent the second major regulatory principle coordinating CESR of *B. subtilis*, LiaRS, BceRS, PsdRS and ApeRS [29]. In addition to these directly CESR-inducible TCSs, the homeostatic TCS WalRK, which coordinates the CW metabolism, and CssRS, which mediates protein secretion stress, are also induced by some triggers of CES.

### Induction of LiaRS by inhibitors of lipid II-cycle

In the present study, the strong induction of the Lia system by bacitracin and vancomycin was confirmed (Fig. 5) [83]. Tunicamycin also activated this system, but to a lesser extent, which is also consistent with a previous study [30]. While all three compounds induced *liaIH*-*liaGFSR*, LiaR was also reported to control two additional targets, the *yhcYZ- yhdA* operon and *ydhE* gene, as suggested by the LiaR-binding sites present upstream of their promoters [53, 84, 85]. The *yhcY* operon was induced weakly by bacitracin (∼2-fold) in this study, whereas *ydhE* did not appear to be responsive to any condition, in line with previous studies [53, 84, 85].

### The response of the detoxification modules controlled by BceRS, PsdRS and ApeRS

In the present study, the *bceAB* operon was specifically and strongly induced by bacitracin (Fig. 5). It was suggested that the BceAB transporter protects the cell by target-protection via transiently freeing lipid II-cycle intermediates from bacitracin [86]. Other lipid II-targeting AMPs, such as the lantibiotics nisin and subtilin, induce the paralogous *psdAB* operon, which is controlled by PsdRS, and responds in a similar way to these two compounds [36]. The *psdAB* operon showed a moderate activation by bacitracin in the present study, which is consistent with previous studies [53, 87] and most likely due to a regulatory cross-activation of *psdAB* expression via BceRS. ApeRS is the least well studied among these three TCSs. So far, the human-derived cationic antimicrobial peptide LL-37 [88], and *Hermetia illucens* larval extract [89] are the only known stimuli of ApeRS. In this study, a ∼2.5 fold induction of its target operon, *apeAB-yxeA*, was observed during penicillin G treatment (Fig. 5). The physiological relevance of this result needs to be further investigated. No other compound was able to activate this TCS.

### The CESR of the secretion stress system CssRS

The CssRS system is part of the quality control mechanisms in protein secretion [37]. The present study demonstrates that the *htrAB* operon was induced by bacitracin, lysozyme, tunicamycin and vancomycin. These results suggest that interfering with membrane-anchored steps of CW biosynthesis also negatively affects protein secretion, thereby generating a stress that is sensed by CssS. Activation of CssRS as part of the CESR of *B. subtilis* has previously also been reported for rhamnolipid treatment [51]. This result demonstrates the close relationship between shuttling CW buildings blocks and proteins to the outside of the cell, indicative for a limited capacity of the membrane for accommodating such export processes, which is an underappreciated of CES that will require further investigations.

### WalRK-dependent CW homeostasis is negatively affected by CES

In the presence of all antibiotics of this study, the genes *cwlO*, *lytE*, *yocH* and *ydjM,* which are positively regulated by WalR [39, 44], exhibited overall downregulated expression (Fig. 6), reflecting the reduction in CW metabolism and suggesting deregulated PG hydrolytic activity when PG synthesis is inhibited. Additionally, *ftsEX*, which is required for CwlO activity [90], was downregulated around five-fold by vancomycin (Table S12).

Conversely, *iseA*, *pdaC* and *sasA,* which are negatively regulated by WalR [39], were released from WalR repression and hence increased in expression (Fig. 6). The strong (30-fold) induction of *iseA* by lysozyme was observed for the first time. Likewise, *pdaC*, which confers lysozyme resistance via de-N-acetylation of PG [91], was also strongly induced by lysozyme. Noteworthy, the response of *pdaC* shows some compound-specificity, in contrast to the almost overall upregulation of *iseA* expression in the presence of all CW inhibitors (Fig. 6). Taken together, the response of WalR-target genes shows that WalRK activity was tuned down to reduce CW metabolism in response to the CES-dependent interference with CW synthesis.

Signature inductions of the CESR in *B. subtilis*

In addition to a comprehensive insight into the regulatory processes, our genome-wide transcriptional profiles on the CESR of *B. subtilis* to different CW antibiotics also unveiled marker genes that were particularly responsive under CES (Fig. 8). Such genes, or rather their target promoters, represent useful candidates for the development of whole-cell biosensors, which are stimulus-specific reporter strains that can be applied for searching for novel antimicrobial compounds [92].

The LiaRS-dependent ***liaIH*** operon has drawn extensive attention because of strong response of up to 1000-fold induction to antimicrobial agents that mostly interfere with lipid II-cycle of PG biosynthesis [30, 53] (Fig. 8). Due to its low basal expression level, the *liaIH* promoter (P*liaI*)-based biosensor is well-established for the identification of cell envelope-active compounds [84, 93].

The genes *yuaF-floT-yuaI*, *yeaA-ydjP*(*-ydjO*), *pbpE-racX*, *pspA-ydjGHI*, *yvlABCD* and *fosB* are under control of σ^W^, which was strongly activated by vancomycin (Fig. 8). The ***yuaF* operon**, was the most sensitive member of the σ^W^ regulon. It is involved in the control of membrane fluidity, which affects CW biosynthesis [94]. The ***yeaA* operon** remains uncharacterized. The first gene of ***pbpE-racX*** operon encodes a penicillin-binding protein PBP4* (endopeptidase), while *racX* codes for an amino acid racemase involved in the production of non-canonical D-amino acids [95]. The first gene of the ***pspA-ydjGHI* operon** encodes the second phage shock protein A (PspA) homolog of *B. subtilis*, in addition to LiaH [84]. YvlC, encoded in the ***yvlABCD* operon**, was identified as a PspC homolog [96]. Finally, ***fosB*** mediates fosfomycin resistance in *B. subtilis* [97].

The expression of the ***yrhH***-***fatR*-*yrhJ*** operon (which is partially involved in fatty acid metabolism [98]) is particularly interesting with regard to its broad spectrum of inducers in this study, including vancomycin; tunicamycin and moenomycin (Fig. 8). This operon is under control of multiple regulators (σ^M^, σ^W^, σ^X^ and FatR) and was assigned into the Group IV during the refinement of ECF σ regulons (Table 2).

In addition to σ^ECF^-dependent genes, the specific induction of the ***ytrABCDEF* operon** by vancomycin was noteworthy. This operon is induced by a narrow range of PG synthesis inhibitors blocking the lipid II precursor, including the glycopeptides vancomycin and ristocetin, which target the terminal D-Ala-D-Ala of the pentapeptide, the glycolipodepsipep-tide ramoplanin, which sequesters lipid II, and plectasin, a fungal defensin that also targets lipid II [67, 99, 100]. Bacitracin was able to activate the *ytr* operon, but to a lesser extent (Fig. 8), consistent with earlier reports [53, 67].

The genes ***htrA*** and *htrB* encode membrane-anchored protein quality control proteases under control of the TCS CssRS. The *htrA* gene was induced by tunicamycin, bacitracin, vancomycin and lysozyme (Fig. 8).

The WalR-dependent gene ***iseA*** was induced by a broad spectrum of antibiotics (Fig. 8). Previously, another member of the WalR regulon, ***sasA***, has been exploited as a biosensor for the discovery of novel CW-active compounds [101, 102]. The present study shows that *iseA* expression was even more sensitive than *sasA* towards cell envelope stress. Therefore, *iseA* may represent a promising candidate for the establishment of reporter strain derived from the WalR regulon.

### Novel genomic features induced by CES

In the course of the resequencing and especially the comprehensive systems biology analysis of *B. subtilis* gene expression [103], numerous novel genomic features were discovered, including non-coding 5’- leader transcripts (5’UTRs), or 3’-extension of genes and operons. Induction of these features is covered by this analysis for the very first time, the results are summarized in Table S21. Most novel features were differentially expressed in only one stress condition, and here the discrepancies between RNAseq and tiling array data is much larger than for the rest of the transcripts. The most consistent of the strongly induced novel features are linked to known regulators of the CESR. This includes ECF-target genes and operons, e.g. S156 (5’UTR of *ddl*, S228 (5’UTR of *yebC*, σ^M^ / σ^V^ regulon), S742 (5’UTR of *yozO*, σ^W^ regulon), S843 (5’UTR of *recU*, σ^M^ regulon) or S659 (downstream of *fosB*). Other novel features with known regulators include S1268 (5’UTR of *htrB*, CssR regulon) and S1275 (3’UTR of the *liaIHGFSR* operon, controlled by LiaR). But the relevance of induction of most of these over 100 novel features by cell wall antibiotics (as listed in Tab. S21) will require further investigations.

### *B. subtilis* biosensors for compound discovery

Based on the signature gene expressions described above, a panel of eleven bioluminescence-based *B. subtilis* biosensors was constructed by transcriptionally fusing the promoters of signature genes to the *luxABCDEF* cassette as a reporter gene (Table S22). We next tested the biosensors for their functionality, sensitivity, response dynamics using vancomycin as a model inducer. A strain with a promoterless *lux* cassette served as a negative control for background luminescence. All biosensors were induced with a dilution series of vancomycin ranging from 0 to 3 µg mL^-1^ (see methods for details). Luminescence and cell density were monitored over time (Fig. S6) and the fold-induction for each condition was calculated (Fig. 9).

**Fig. 9.**
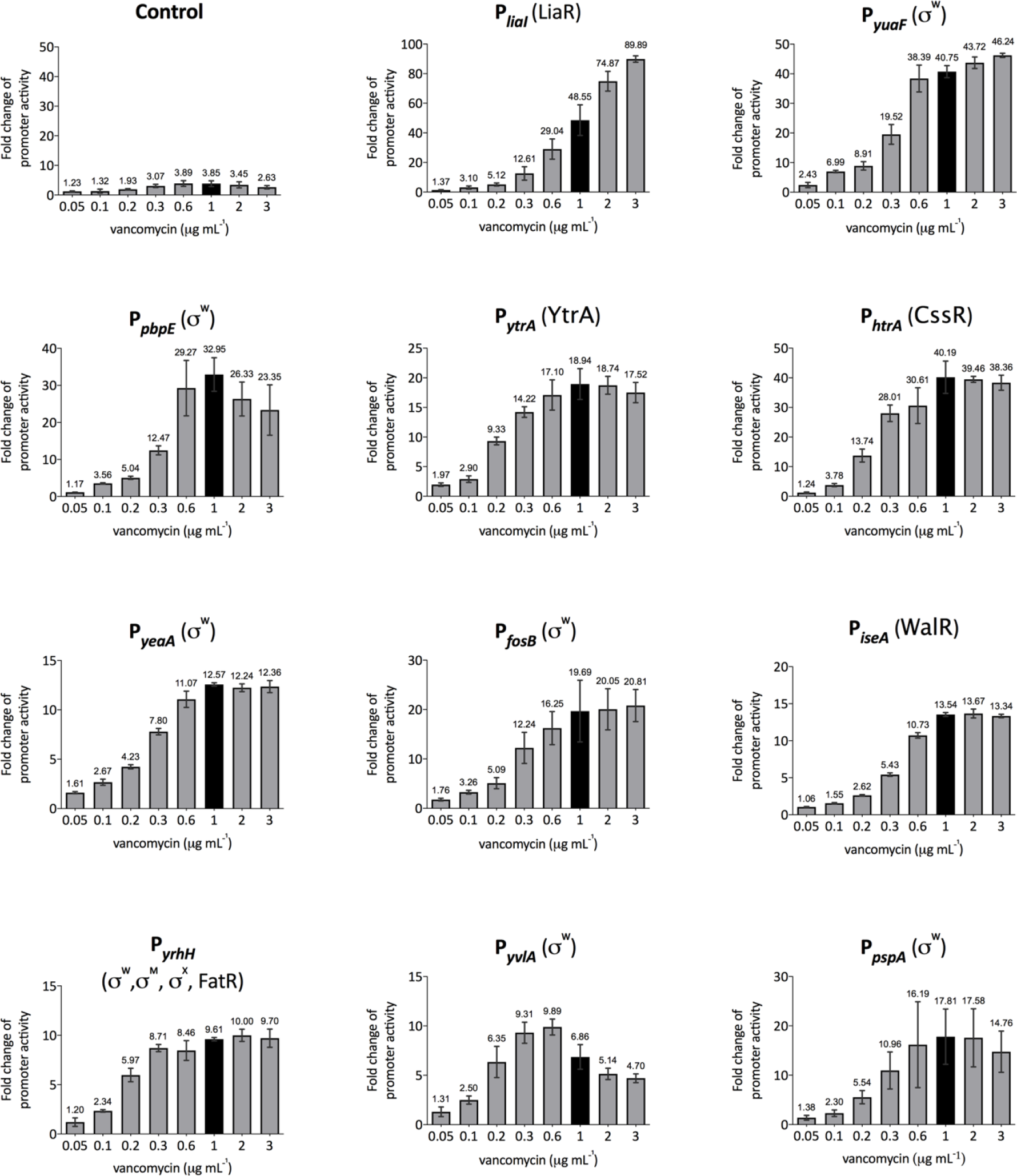
Dose-dependent response of *B. subtilis* biosensors to vancomycin. The strain harboring promoterless-*lux* fusion was used as negative control. The biosensors after one hour of growth in a plate reader were treated with a series of concentrations of vancomycin, respectively, as indicated on the y-axis of each graph. The fold-change of promoter activity was obtained by normalizing the highest promoter activity, represented by luciferase activity (RLU/OD600, as shown in Fig.17 and Fig. S3), after the addition of vancomycin to the promoter activity under untreated condition (0 µg mL^-1^) at the same time point. The promoter activities under 1 µg mL^-1^ of vancomycin treatment are highlighted in black in order to compare with RNAseq data. Error bars indicate the standard deviation of at least three biological replicates.

All biosensors exhibited a dose-dependent response to vancomycin and a good correlation – or even a higher dynamic range – was observed for the biosensors relative to the transcriptome data. e.g. the P*htrA* biosensor was activated approx. 40-fold, while only a five-fold induction of *htrA* expression was observed by RNAseq. Likewise, the P*pbpE* biosensor was induced 33-fold, while only a nine-fold induction of *pbpE* expression was detected. In both cases, this was due to the induction kinetics, since P*htrA* and P*pbpE* responded more slowly to vancomycin than other promoters (Fig. S6). Consequently, the highest induction value chosen for the biosensor assays occurred at 30-45 minutes post-induction, while the induction levels of *htrA* and *pbpE* in RNAseq were determined after 10 min of induction.

No major differences were observed between the sensitivity of the promoters to vancomycin. The threshold concentration for P*yuaF* and P*ytrA* induction was 0.05 µg mL^-1^, while all other promoters starting responding at 0.1 µg mL^-1^ (Fig. 9 and Fig. S6). The maximum activity of most promoters occurred at 1 µg mL^-1^ of vancomycin, while higher concentrations did neither increase nor decrease the promoter activity further. Only P*liaI* activity was still increasing at 2 or even 3 µg mL^-1^ of vancomycin. Conversely, P*yvlA* exhibited the highest activity at 0.6 µg mL^-1^, followed by declined activity at higher concentration treatments.

Noteworthy, P*iseA* – but not any other promoters – displayed increased activity at late-exponential phase, even in the absence of stimulus (Fig. S6), due to a negative regulation by WalRK. During exponential phase, WalRK is highly active, thereby repressing *iseA* expression. With a decrease of CW growth at the onset of the stationary phase, WalRK activity is also declining, thereby releasing *iseA* from WalR repression. The P*iseA* biosensor was also tested with penicillin G, and similar dose-response behavior was observed (Fig. S7).

This novel panel of *B. subtilis* biosensors expands the range of biosensors already available for CW antibiotics. Their functionality and performance were verified based on their dose-response behavior to vancomycin, but some additionally responded to other antibiotics such as bacitracin and moenomycin (Fig. S8), underscoring their potential to detect new cell envelope-active compounds to facilitate the discovery of novel antimicrobial compounds.

## Discussion

Bacteria living in complex environments, like the soil, compete with numerous other species for ecological niches and the scarce nutrients. Biological warfare – in the form of antibiotics – is one aspect of this competition, which allows bacteria to strive and prosper in the face of competitors. PG synthesis is a prime target for many antibiotics, due to its crucial role, and cells need to continuously monitor its integrity, in order to mount swift responses in the case of envelope damage. For over two decades, the underlying regulatory network of the CESR has been thoroughly studied in the Gram-positive soil bacterium *B. subtilis* [24, 104], but a systematic and comparative analysis has been missing so far. Here, we presented the results of such a comprehensive and highly standardized transcriptomic profiling study. We chose eight antimicrobials, including seven CW antibiotics that inhibit PG synthesis from early cytoplasmic steps (fosfomycin, D-cycloserine) via the membrane anchored lipid II cycle (tunicamycin, bacitracin, vancomycin) to the extracellular polymerization steps (penicillin G, moenomycin) and lysozyme, which degrades the existing CW (Fig. 1). Highly standardized experimental conditions (inhibitory but sublethal antibiotic concentrations and only 10 min induction) were chosen to exclusively monitor the primary, that is, initial antibiotic-specific responses. Moreover, two independent state-of-the-art transcriptomic technologies, RNAseq and the latest generation of tiling array, were applied in parallel to solidify the data and exclude technical biases. This approach enabled us gaining a comprehensive picture of the CESR of *B. subtilis*, providing an unsurpassed resolution on how an organism perceive threats and damages to its envelope. By mapping the transcripts on the updated genome sequence, we were also able to identify novel features, such as non-coding RNAs, even for stimulons that had been thoroughly analyzed in the past, such as the vancomycin or bacitracin stress responses. Taken together, our work provides a comprehensive reference analysis for future studies on the CESR in *Bacillus* species and related Firmicutes bacteria.

Applying two independent technologies for monitoring transcriptome profiles enabled us to both validate the data and also identify potential technical biases in our analysis. While RNAseq is currently the most widely used standard for monitoring genome wide transcriptional profiles, high-resolution *B. subtilis* tiling arrays were thoroughly evaluated in the largest systems biology study for this organism [103], and served as an internal standard for our work. Overall, both approaches resulted in highly comparable datasets (Figs. 4, 5, 6, 8), demonstrating the robustness of transcriptomic studies, irrespective of the specific technology applied to quantify the transcripts. In contrast, the experimental conditions applied for cultivating the cells are highly critical, as shown by the differences found in the vancomycin stimulons when the antibiotic concentration varied by only a factor of four (0.25 vs. 1 µg/ml vancomycin). While the overall pattern was comparable, the higher antibiotic concentration resulted in higher fold-induction of the target genes and also in the additional induction of secondary, less specific responses, such as the SigB-dependent general stress response (Fig. 7).

Remarkably, applying standardized and comparable conditions to seven antibiotics that interfere with successive steps of PG synthesis also highlighted how the resulting stresses are perceived by the regulatory systems involved in orchestrating the CESR in *B. subtilis*. While a strong and differentiated response was observed for all compounds interfering with membrane-associated steps of PG synthesis (tunicamycin, bacitracin, vancomycin), for the compounds inhibiting either cytoplasmic (fosfomycin, D-cycloserine) or extracellular steps (moenomycin, penicillin G) no clear primary transcriptional signatures were observed and none of the known CESR systems was significantly triggered (Figs. 2-6). The strong and differential response to antibiotics interfering with membrane-anchored steps of PG synthesis is well documented [51–53]. It highlights the key role of the lipid II cycle as the bottleneck process in PG synthesis. By blocking this process, PG synthesis becomes dramatically dysbalanced: while soluble PG precursors accumulate on the cytoplasmic side, the growing murein sacculus is depleted for building blocks, and the action of PG hydrolases, which precedes but is normally well-coordinated with the incorporation of new material, gets deregulated and weakens the CW further. In addition to the almost instant effect that blocking the lipid II cycle has on PG biosynthesis, the site of this inhibition at the membrane is also ideally suited for perception by the membrane-anchored sensors of CES, e.g. the sensor kinases and the anti-σ factors (or the proteases degrading them). Our transcriptional profiles strongly suggest that the CESR of *B. subtilis* has specifically evolved to immediately perceive interference with the lipid II cycle as the most reliable indicator of future CW damage. This hypothesis is in agreement with numerous additional transcriptomic studies from both Gram-positive and Gram-negative bacteria that draw a similar picture [51].

In contrast, interfering with cytoplasmic or extracytoplasmic steps is hardly detected by the cells at first, at least under the chosen experimental conditions. In the case of fosfomycin and D-cycloserine, the pool of soluble CW precursors was most likely not yet depleted after 10 min to significantly affect the successive steps, such as the lipid II cycle, and elicit a stronger transcriptional response. While using longer times between antibiotic induction and cell harvest might have resulted in a stronger and clearer transcriptional profile, it was not the aim of this study to monitor downstream transcriptional effects of metabolic depletion, but rather to provide a comprehensive picture of the primary CESR of *B. subtilis*. Towards this end, our data clearly indicates that the initial, cytoplasmic reactions do not represent suitable stimuli to provide the cell with a sensitive read-out for CES. The poor primary (within 10 min) response to the extracellular inhibitors of CW biosynthesis moenomycin and penicillin, which was also observed for other bacteria [51], might result from the plasticity of the PG meshwork to adapt to changing, often challenging, environmental conditions, favored by the multiplicity and often redundancy of the main players. TP and TG reactions are tightly coordinated between them and with the action of PG hydrolases, in order to incorporate new building blocks in the dynamically growing PG network [105]. In rod-shaped bacteria this process is coordinated by combining the necessary enzymes in highly motile PG biosynthetic complexes that are organized by cytoskeletal elements. Sidewall elongation is effected by the Rod complex, associated to the actin homolog MreB, and septum formation is effected by the divisome, associated to the tubulin homolog FtsZ [105, 106]. In *B. subtilis*, PG synthesis is additionally mediated by aPBPs (bifunctional PBPs with both TG and TP activity) functioning outside these complexes [107]. Moenomycin targets the glycosyltransferase activity of aPBPs [20] but not of the essential SEDS glycosyltransferases RodA and FtsW that are associated to the Rod complex and to the divisome, respectively. In agreement with this, in *B. subtilis* aPBPs are not essential and PG synthesis continues in moenomycin-treated cells [108]. Thus, the absence of a rapid transcriptional response when sublethal concentrations of moenomycin are added to exponentially growing cells is not too surprising. In contrast, penicillin blocks the TP activity of PBPs, which includes TP by the bifunctional aPBPs and by the monofunctional bPBPs associated to RodA and FtsW, and thus all TP activity in the sacculus. However, the effect of sublethal concentrations of penicillin in sacculus crosslinking within 10 min may not be sufficient to trigger a CESR, or else be compensated by reducing the activity of PG hydrolases. Bacterial cells are known to be able to accommodate variations in the amount, the fine composition or the crosslinking of PG, which has been proposed to help to deal with transient inhibitions of PG synthesis.

Finally, lysozyme hydrolyzes the glycosidic linkage between GlcNAc and MurNAc, which can rapidly compromise the integrity of the sacculus and result in cell lysis. Furthermore, lysozyme directly binds to the membrane-anchored anti-σ^V^ factor, RsiV, directly inducing σ^V^ activation and thus the expression of proteins required for lyzozyme resistance [28]. The response to lysozyme is therefore rapidly detected by CESR systems, but compound-specific rather than a CW damage-triggered response. Noteworthy, lysozyme and vancomycin show almost inverted induction/repression patterns in their transcriptional profiles (Fig. 2). The reason for this odd behavior remains to be investigated.

## Conclusion

Our comprehensive survey of the primary CESR of *B. subtilis* demonstrates that monitoring (the inhibition of) the lipid II cycle is the primary check point to monitor the state of PG biosynthesis and orchestrate adequate countermeasures before lethal damage can occur to the envelope, as has been thoroughly demonstrated in case of the bacitracin stress response [24, 63, 109]. Our work not only provides a future reference point for the global transcriptional CESR, it also serves as a direct comparison of the performance of two profiling approaches – RNAseq vs. tiling arrays – and provides a collection of highly sensitive whole cell biosensors for monitoring CESR. Such biosensors could be useful tools in the antibacterial research field. At a time when the spread of bacterial resistance has become a global threat, the PG cell wall, an essential bacterial structure lacking in higher organisms, remains the most prominent target for antibacterial therapy [4].

## Materials and Methods

### Strains and growth conditions

*Bacillus subtilis* BaSysBio wild type (Nr. 92 in AG Mascher *Bacillus* collection) was used for transcriptomic study, and routinely grown in Lysogeny Broth (L3522-LB broth, Sigma-Aldrich) (tryptone, 10 g L^-1^; yeast extract, 5 g L^-1^; NaCl, 10 g L^-1^) at 37 °C with aeration. *B. subtilis* biosensors were derived from *B. subtilis* W168 (Table S22). *B. subtilis* W168 strains and *E. coli* were routinely cultivated in LB (Luria/Miller, Carl Roth) (tryptone, 10 g L^-1^; yeast extract, 5 g L^-1^; NaCl, 10 g L^-1^) at 37 °C with aeration. Solid media contained 1.5% (w/v) agar. Selective media for *B. subtilis* W168 contained chloramphenicol (5 µg mL^-1^), and for *E. coli* contained ampicillin (100 µg mL^-1^).

### DNA manipulation and plasmid construction

General cloning procedure, such as PCR, restriction enzyme digestion and ligation, was performed with enzymes and buffers from New England Biolabs^®^ (NEB, Ipswich, MA, USA) according to respective protocols. Q5^®^ High-Fidelity DNA polymerase was used for PCRs in case the resulting fragment was further used, otherwise OneTaq^®^ was the polymerase of choice. PCR purification was performed using the Hi Yield^®^ PCR Gel Extraction/PCR Clean-up Kit (Süd-Laborbedarf GmbH (SLG), Gauting, Germany). Plasmid preparation was performed using the Hi Yield® Plasmid Mini-kit. The resulting constructs were verified by sequencing.

To generate promoter-*lux* fusions, the promoters were amplified from the genomic DNA of *B. subtilis* using respective primer pairs (Table S2424) and cloned into pBS3C*lux*, a reporter vector in the *B. subtilis* BioBrick Box [110]. The vector and the details of plasmid construction are described in Table S2323.

### E. coli and B. subtilis transformation

The chemically competent *E. coli* cells were used for cloning. *E. coli* transformation was done as: 50 µL of *E. coli* competent cells were thawed on ice for about 10 min; ½ (or the whole) volume of ligation reaction mix was added to the cells and mixed gently; After 30 min incubation of the tube on ice, the cells were heat-shocked for 45 seconds at 42 °C and placed back on ice immediately for at least 2 min. 900 µL LB medium was added to the tube and incubated at 37 °C for 1 hour with shaking; 50 µL or 100 µL (depending on experiments) of the recovery culture were plated on selective LB plates and incubated at 37 °C overnight.

*B. subtilis* transformation was performed as: 10 mL MNGE medium was inoculated 1:100 from overnight cultures of the recipient *B. subtilis* strain. Cultures were grown to OD600 of 1.1-1.3 at 37 °C, 200 rpm; 400 µL of the cells were taken into sterile glass tube for transformation and DNA was added (2 µg linearized plasmid DNA). The mixture was incubated for 1 h at 37 °C with agitation and then 100 µL expression mix were added. After another 1 hour of incubation at 37 °C with agitation, 50 µL or 100 µL (depending on experiments) of the culture were plated on selective LB plates and incubated at 37 °C overnight. Successful integration of fragment into *B. subtilis* genome was confirmed via colony PCR. MNGE medium: 9.2 mL 1X MN medium (136 g L^-1^ dipotassium phosphate x 3 H2O, 60 g L^-1^ monopotassium phosphate, and 10 g L^-1^ sodium citrate x 2 H2O), 1 mL glucose (20%, w/v), 50 µL potassium glutamate (40%, w/v), 50 µL ammonium ferric citrate (2.2 mg mL^-1^), 100 µL tryptophan (5 mg mL^-1^), and 30 µL magnesium sulfate (1 M). Expression mix: 500 µL yeast extract (5%, w/v), 250 µL casamino acids (10%, w/v), 50 µL tryptophan (5 mg mL^-1^) and 250 µL H2O.

### Sample preparation and RNA isolation for RNAseq analyses

The sublethal concentration of the compounds against *B. subtilis* was firstly determined prior to the induction experiment. The overnight culture was made from fresh single colony of *B. subtilis* grown at 37 °C overnight in a shaker at 200 rpm. 10 mL LB medium in a 100 mL flask was inoculated with overnight culture by the ratio of 1:100 and incubated at 37 °C with agitation until OD600 of around 0.4-0.5 as Day Culture 1. Next, in a 2 L flask, 200 mL LB was inoculated with Day Culture 1 to OD600 of 0.01 and incubated at 37 °C in a shaker with measurement of OD600 every 30 min until it reached to ∼0.4 as Day Culture 2. Subsequently, Day Culture 2 was split into fractions of 25 mL in 250 mL flasks, which were induced with different concentrations of the compounds, leaving one un-induced as control. OD600 of each fraction was measured every 30 min up to 2 hours. The concentrations that inhibit *B. subtilis* growth as shown in Fig. S1 (the growth curve at 1 µg mL^-1^) were determined as sublethal concentrations and further applied to induce *B. subtilis* in the following procedure.

To prepare bacterial cell samples for RNA isolation, Day Culture 1 and 2 were prepared as described above. The Day Culture 2 at OD600 of ∼0.4 were split into 25 mL fractions in each pre-warmed 250 mL flask with an appropriate amount of compounds added already. The cultures were then immediately incubated at 37 °C, 220 rpm. After exact 10 min of induction, the cultures were immediately transferred into accordingly labeled 50 mL centrifugation tubes (Sarstedt^TM^, Thermo Fisher Scientific) and put into ice/NaCl bath (ice: NaCl, 3: 1 (v/v)) to efficiently terminate the induction reaction. Afterwards, the cultures were centrifuged in a precooled centrifuge at 4 °C, 8000 rcf for 2-3 min. The supernatant was directly decanted from the culture. Cell pellets were then snap-frozen in liquid nitrogen and stored at -80 °C until use. Every treatment and control samples were made in triplicate. To isolate total RNA, the *B. subtilis* cell pellets were re-suspended in 200 µL killing buffer (20 mM Tris/HCl pH 7.5, 5 mM MgCl2, 20 mM NaN3) and transferred to pre-frozen (in liquid nitrogen) homogenizer vessel including the steel ball, followed with disruption in a homogenizer (Mikro-Dismembrator S, Sartorius, Germany) for 2 min at 2600 rom. The cell powder was re-suspended in 4 mL pre-warmed lysis buffer (116.16 g GTC, 2.05 mL sodium acetate (3 M pH 5.2, final conc. 0,025 M), 12.5 mL lauroylsarcosine (10%, final conc. 0.5%), add DEPC-treated H2O to 250 mL) and transferred into four 2 mL reaction tubes with 1 mL in each.

1 mL Phenol Mix (Phenol: Chloroform: Isoamylalcohol 25: 24: 1, pH 4.5-5, ROTI^®^ Aqua-P/C/I, for RNA extraction, Carl Roth, Germany) was added to 1 mL of lysed cells, followed with extraction for 5 min by vigorous mixing using multi-vortex (Eppendorf). The mixture was centrifuged for 5 min at 12,000 rcf. Afterwards, around 800 µL supernatant were transferred into a fresh 2 mL reaction tube with 800 µL Phenol Mix added. A second extraction followed with centrifugation was conducted. Around 700 µL supernatant were transferred into a fresh 2 mL reaction tube with addition of the same volume of Chloroform Mix (Chloroform: Isoamylalcohol 24: 1, Roti^®^-C/I, for nuclear acid extraction, Carl Roth, Germany). The mixture was extracted and then centrifuged as before. Around 500 µL of the supernatant were transferred afterwards into a fresh 2 mL reaction tube, followed by the addition of 50 µL (1/10 volume) sodium acetate (3 M, pH 5.2) and 1 mL (2 volume) isopropanol. The mixture was mixed by inverting and incubate at -80°C overnight. Next day, the precipitation was centrifuged for 30 min at 15,000 rcf, 4 °C in a precooled centrifuge. The supernatant was removed afterwards, and the pellet was washed twice with 1 mL of 70% ethanol, followed by centrifugation for 5 min at 15,000 rcf, 22 °C. Then, the supernatant was decanted directly and the pellet was dried for about 10 min at room temperature. After that, the RNA pellet was dissolved in 20-50 µL DEPC treated H2O. Two RNA pellets from one sample were at the end combined into one tube and stored at -80°C until further use.

### Sample preparation and RNA isolation for tiling array analyses

Overnight cultures of *Bacillus subtilis* wild type strain (grown at 30°C, 200 rpm shaking) were diluted in fresh medium to OD600nm 0.01. Cells were grown at 37°C to mid-exponential phase (OD600nm 0.4-0.5), re-diluted to OD600nm 0.01 and further grown at 37°C, 200 rpm until mid-exponential phase.

Cultures grown to OD600nm 0.4-0.5 were split in 100 mL aliquots for induction with sub-lethal concentrations of antibiotics, leaving one fraction as uninduced control. After 10 min of incubation at 37°C, 35 mL of culture were mixed with 15 mL of ice-cold killing buffer (20 mM Tris/HCl [pH 7.5], 5 mM MgCl2, 20 mM NaN3) and immediately centrifuged (5 min, 6000 rpm 4°C). Pellets were frozen in liquid nitrogen and kept at -80°C.

RNA samples were prepared as in Nicolas et al., 2012 with only slight modifications [56]. Briefly, cells were mechanically lysed by bead beating (Mikro-Dismembrator S from Sartorius) as described previously. For RNA extraction, 1 volume of acid phenol (Roti-Aqua-phenol from Carl Roth) was mixed (5 min, 1400 rpm) with 1 volume of cell lysate. Three rounds of extraction with chloroform/isoamyl- alcohol 24:1 in Tris-HCl [pH 8] were performed before RNA precipitation with 3M sodium acetate and isopropanol overnight at -20°C. RNA was collected by centrifugation (15000 rpm, 15 min, 4°C), washed with 70% ethanol, resuspended in 75 µL ddH2O and digested with DNaseI (QIAGen RNase-Free DNase set (ref. n°79524) for 10 min at RT. Samples were cleaned-up using the Norgen Concentration Micro Kit (ref. n°23600) according to manufacturer instructions. RNA concentration was determined by Nanodrop and RNA quality using Agilent Bioanalyzer chip. Hybridization on tiling array chips were realized at PartnerChip.

### Tiling array analysis

Analysis was realized by pooling results from three experiments of each condition. Tiling array data are obtained with a strand-specific resolution of 22 bp in the different conditions considered in this study. The analysis used the signal processing and gene-level aggregation procedures used in (P. Nicolas et al. 2012).

Statistical comparison of the 3 biological replicates for each of the considered conditions relied on the functions “lmFit” and “eBayes” of R package “limma” (Smyth 2004). Control of the False Discovery Rate relied on q-values obtained with R package “fdrtool” (Strimmer 2008), where the p-values in input are from “eBayes”. Genes were then considered as differentially expressed if q-values were at least less than 0.05.

### RNA sequencing and analysis

The RNA library quality was verified using LabChip GX Touch HT Nucleic Acid Analyzer. rRNA was subtracted from the samples with the Illumina Ribo-Zero rRNA removal Kit (Bacteria) according to manufacturer instructions. The cDNA library was prepared using the NEB Ultra directional RNA library prep kit for Illumina according to instructions and sequencing was performed on an Illumina HiSeq3000 system. For analysis, the quality of the raw sequencing files was verified using MultiQC. Next, the sequences were aligned to the *Bacillus subtilis* subsp. subtilis str. 168 complete genome (NC_000964.3) using Bowtie 2 (Bowtie 2: 2.4.1). Unmapped reads were filtered with Samtools (Samtools: 1.10). Mapped reads were sorted, and converted to bam file with Samtools (Samtools: 1.10). Gene counts of aligned reads were quantified using FeatureCounts. The counts were normalized using DESeq2 and a differential gene expression was calculated (DESeq2: 1.28.0, r-base: 4.0.2). The DESeq 2 comparisons were combined and enriched to an Excel sheet using in-house scripts. Non-treatment condition was used as the reference point. Genome annotation (in GFF format) was gathered from BSGatlas (Version 1.0). All raw sequencing data, the processed data files and differential expression data are deposited at GEO (Gene Expression Omnibus) platform with accession number GSE160345.

The hierarchical clustering was performed using Heatmap.2 with in-house scripts and the clustering was based on the log2 fold-change value of the 327 genes. The graphs shown in the text were generated using GraphPad.

### Luciferase assay

The luciferase activity of *B. subtilis* reporter strains carrying *luxABCDE* operon was assayed using a Synergy™ NEO multi-mode microplate reader from BioTek^®^ (Winooski, VT, USA). The reader was controlled by the software Gen5™ (Version 2.06). Luminescence assays were carried out as followed: 10 mL LB medium (w/o antibiotics) were inoculated 1:1000 from overnight cultures (grown with respective antibiotics) and grown to OD600 of 0.2-0.3. Then, day cultures were diluted to an OD600 of 0.01 and 200 µL were transferred into wells of 96-well plate (black wall, clear bottom; Greiner Bio-One, Frickenhausen, Germany). After one hour of incubation, 5 µL of vancomycin with corresponding concentrations were added to the culture, respectively. Non-treatment was added with the same amount of sterile water as the control. The program was set up for incubation of the plate at 37 °C with agitation (intensity: medium) and the OD600 as well as the luminescence was recorded every 5 min for at least 18 hours. Luciferase activity (RLU/OD600) was defined as the raw luminescence output (relative luminescence units, RLU) normalized to OD600 corrected by medium blank at each time point.

## Acknowledgments

Work in the Carballido-Lόpez laboratory was funded by the European Research Council (ERC) under the European Union’s Seventh Framework Program (FP7) and the Horizon 2020 research and innovation program (grant agreement No 311231 and grant agreement No 772178, respectively, to R.C.-L.). Work in the Mascher laboratory was supported by grants from the Deutsche Forschungsgemeinschaft (DFG grant MA2837/3-2) in the framework of the priority program SPP1617 ‘Phenotypic Heterogeneity and Sociobiology of Bacterial Populations’ and the Bundesministerium für Bildung und Forschung (BMBF) in the framework of the ERAnet Synthetic Biology (project ERASynBio2- ECFexpress). Q.Z. was supported by the China Scholarship Council and the Graduate Academy of Technische Universität Dresden.

## Author contributions

T.M and R.C.-L. conceived the study. Q.Z., C.C. and P.F. performed experiments. Q.Z. and C.C. analyzed the data and generated all figures and tables. C.G., D.M., R.R.M. and V.F. were involved in RNA isolation and/or analyzing the RNAseq and tiling array experiments. D.W. supervised the experimental work in the Mascher group. Q.Z. and T.M. wrote the original draft of the manuscript. All authors took part in the manuscript revision.

## List of figures, tables and supplemental material

Fig. 1 The peptidoglycan biosynthetic pathway showing sites of action of the compounds applied in the study

Fig. 2 Hierarchical clustering analysis of the transcriptional profiles of B. subtilis in response to cell envelope-active compounds.

Fig. 3 Graphic presentation of the stimulons.

Fig. 4 Induction profiles of sigW, sigM, sigV and sigX by the CESs.

Fig. 5 Transcriptional response of two-component systems under cell envelope stresses.

Fig. 6 Transcriptional response of WalR regulon to cell envelope-active compounds.

Fig. 7 Vancomycin stimulon (RNAseq vs Tilling array).

Fig. 8 Expression patterns of CESR marker genes.

Fig. 9 Dose-dependent response of B. subtilis biosensors to vancomycin.

Fig. S1 Effect of treatment of different concentrations of vancomycin on B. subtilis growth.

Fig. S2 Transcriptional response of rRNA-coding gene to different CESs.

Fig. S3 Hierarchical clustering analysis of the transcriptional profiles of B. subtilis in response to cell envelope-active compounds (tiling array).

Fig. S4 Clustering analysis of the ECF sigma regulons.

Fig. S5 Promoter consensus sequences.

Fig. S6 Dose-response of B. subtilis biosensors to vancomycin.

Fig. S7 Dose-response of B. subtilis biosensor with promoter PiseA to penicillin G.

Fig. S8 Induction of B. subtilis biosensors to bacitracin and/or moenomycin.

Table 1 Top 5 differentially expressed genes in each of the stimulons.

Table 2 Overview of ECF σM, σW, σX and σV regulons (RNAseq & Tiling array).

Table S1 Complete table of differentially expressed genes with RNAseq. Table S2 Complete table of differentially expressed genes with tiling array.

Table S3 The full list of genes applied to generate heatmap with the corresponding clusters assigned marked.

Table S4 Genes induced or repressed (≥2 fold) in fosfomycin stimulon with RNAseq.

Table S5 Genes induced or repressed (≥2 fold) in fosfomycin stimulon with tiling array.

Table S6 Genes induced or repressed (≥2 fold) in D-cycloserine stimulon with RNAseq.

Table S7 Genes induced or repressed (≥2 fold) in D-cycloserine stimulon with tiling array.

Table S8 Genes induced or repressed (≥5 fold) in tunicamycin stimulon with RNAseq.

Table S9 Genes induced or repressed (≥5 fold) in tunicamycin stimulon with tiling array.

Table S10 Genes induced or repressed (≥5 fold) in bacitracin stimulon with RNAseq.

Table S11 Genes induced or repressed (≥5 fold) in bacitracin stimulon with tiling array.

Table S12 Genes induced or repressed (≥5 fold) in vancomycin stimulon with RNAseq.

Table S13 Genes induced or repressed (≥5 fold) in vancomycin stimulon with tiling array.

Table S14 Genes induced or repressed (≥2 fold) in moenomycin stimulon with RNAseq.

Table S15 Genes induced or repressed (≥2 fold) in moenomycin stimulon with tiling array.

Table S16 Genes induced or repressed (≥2 fold) in penicillin G stimulon with RNAseq.

Table S17 Genes induced or repressed (≥2 fold) in penicillin G stimulon with tiling array.

Table S18 Genes induced or repressed (≥5 fold) in lysozyme stimulon with RNAseq.

Table S19 Genes induced or repressed (≥5 fold) in lysozyme stimulon with tiling array.

Table S20 Genes for clustering analysis of ECF sigma regulons.

Table S21 New features significant in at least one condition of RNAseq and tiling array.

Table S22 Bacterial strains used in this study.

Table S23 Vector and plasmids used in this study.

Table S24 Oligonucleotides used in this study.

---

**Fig. S1.**
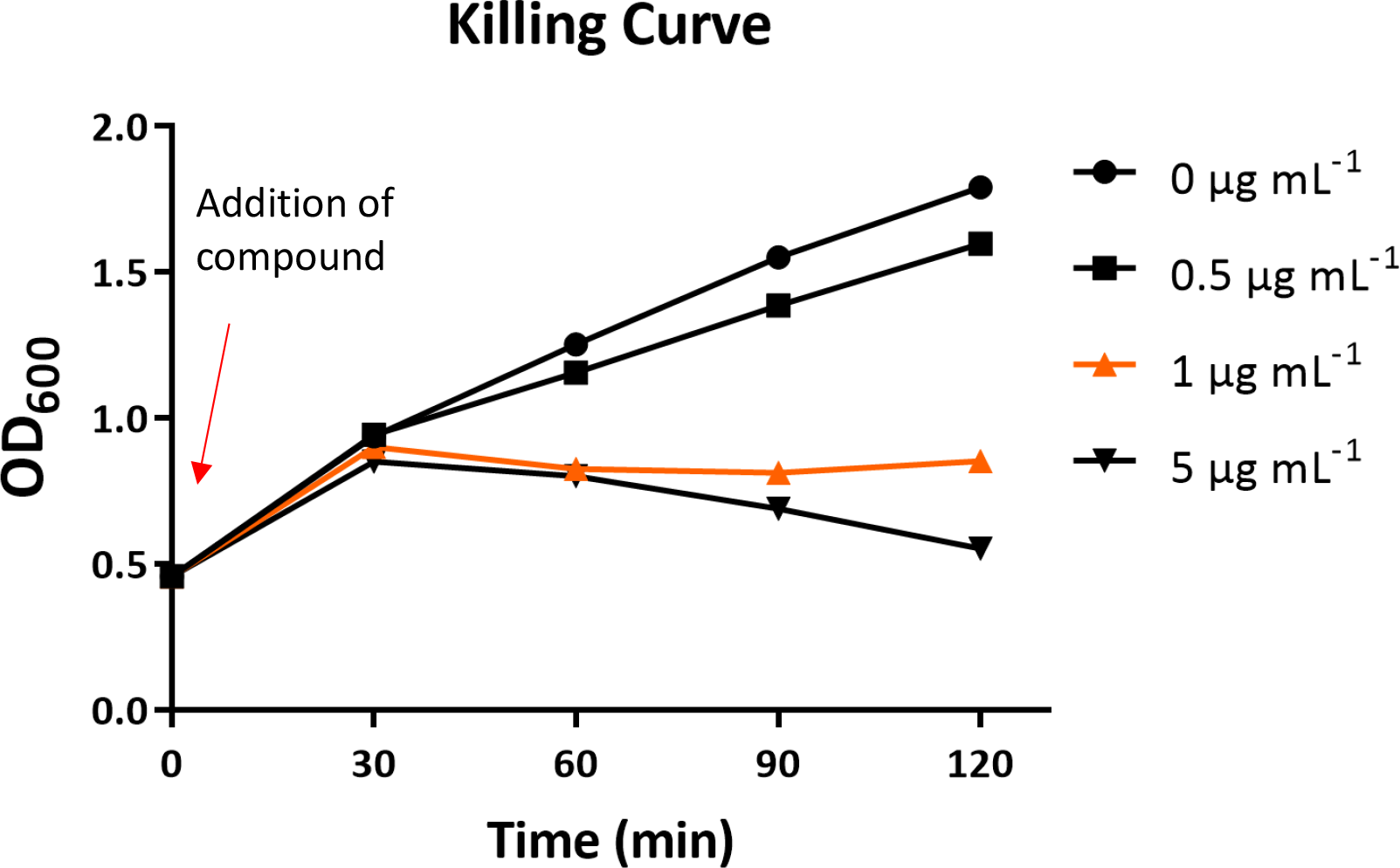
Effect of treatment of different concentrations of vancomycin on *B. subtilis* growth. The killing curve assay results for vancomycin in RNAseq experiment was represented here to depict the definition of the sublethal concentration applied in this study. The sublethal concentration (1 μg mL^-1^) for vancomycin is highlighted in orange. The determination of the concentration for other compounds followed the same fashion.

**Fig. S2.**
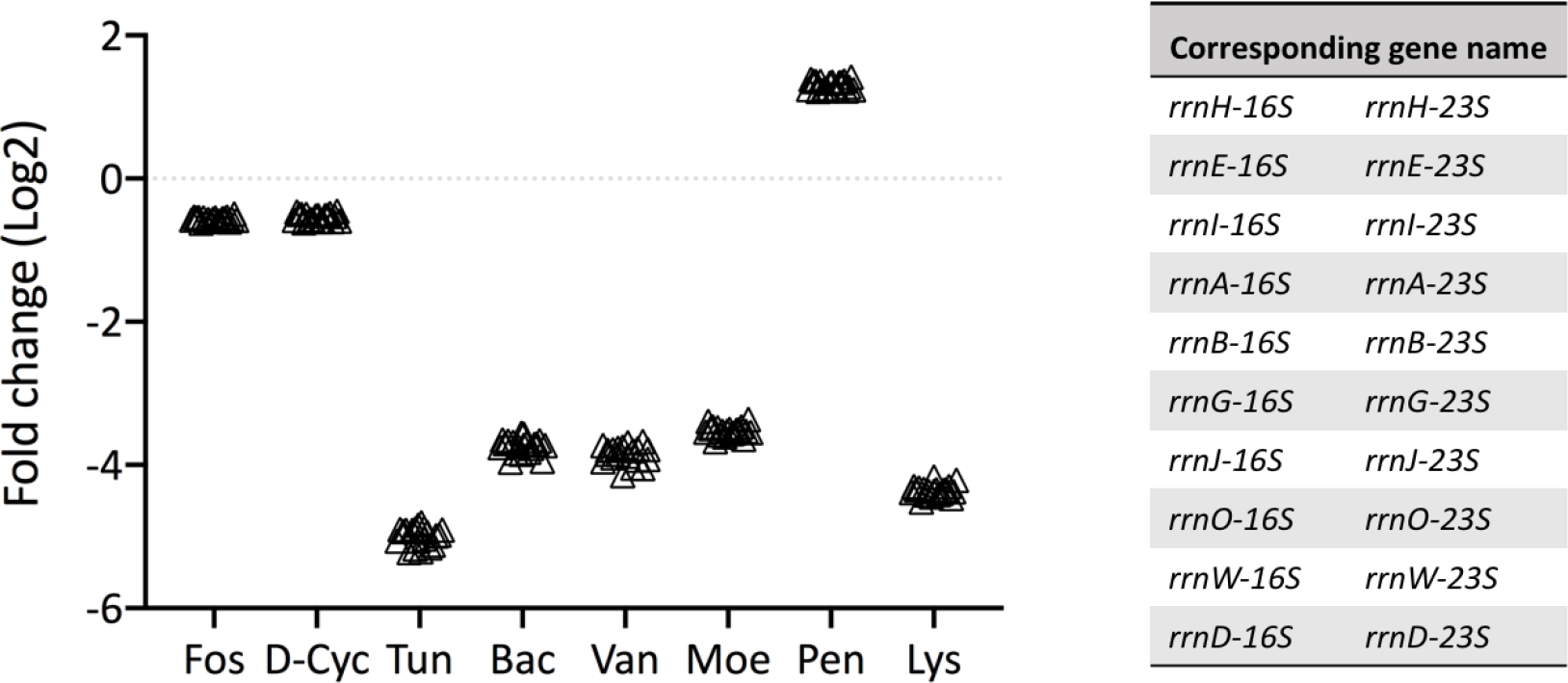
Transcriptional response of rRNA-coding gene to different CESs. The 20 rRNA-coding genes are listed in the table on the right. The log2 fold change of these genes is displayed in the graph on the left with each gene represented by an open triangle. The triangles in each stimulon overlap because of their similar differential gene expression.

**Fig. S3.**
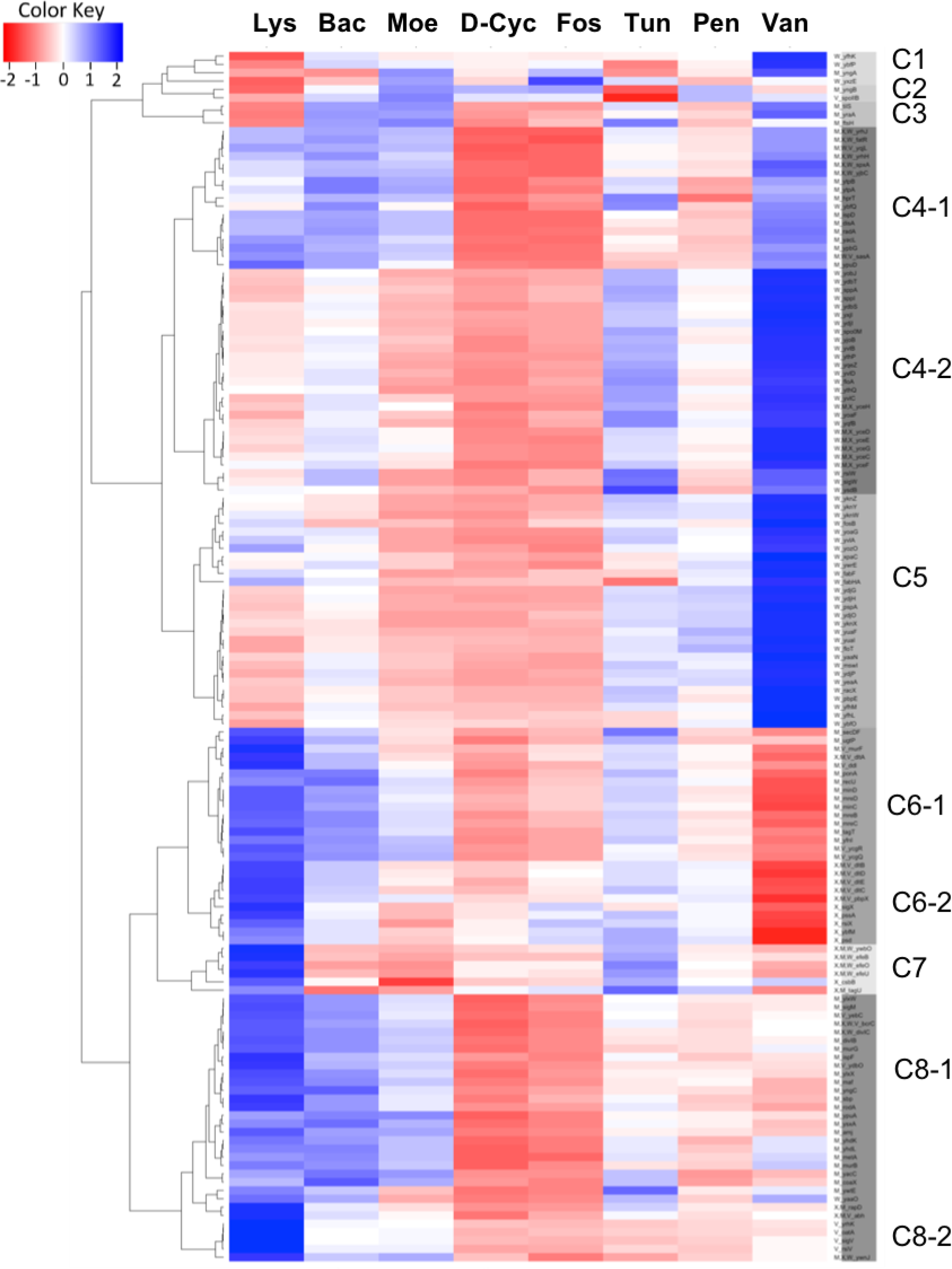
Heatmap (tiling array) Hierarchical clustering analysis of the transcriptional profiles of *B. subtilis* in response to cell envelope-active compounds (tiling array). Tiling array data in log2 was used to generate the heatmap.

**Fig. S4.**
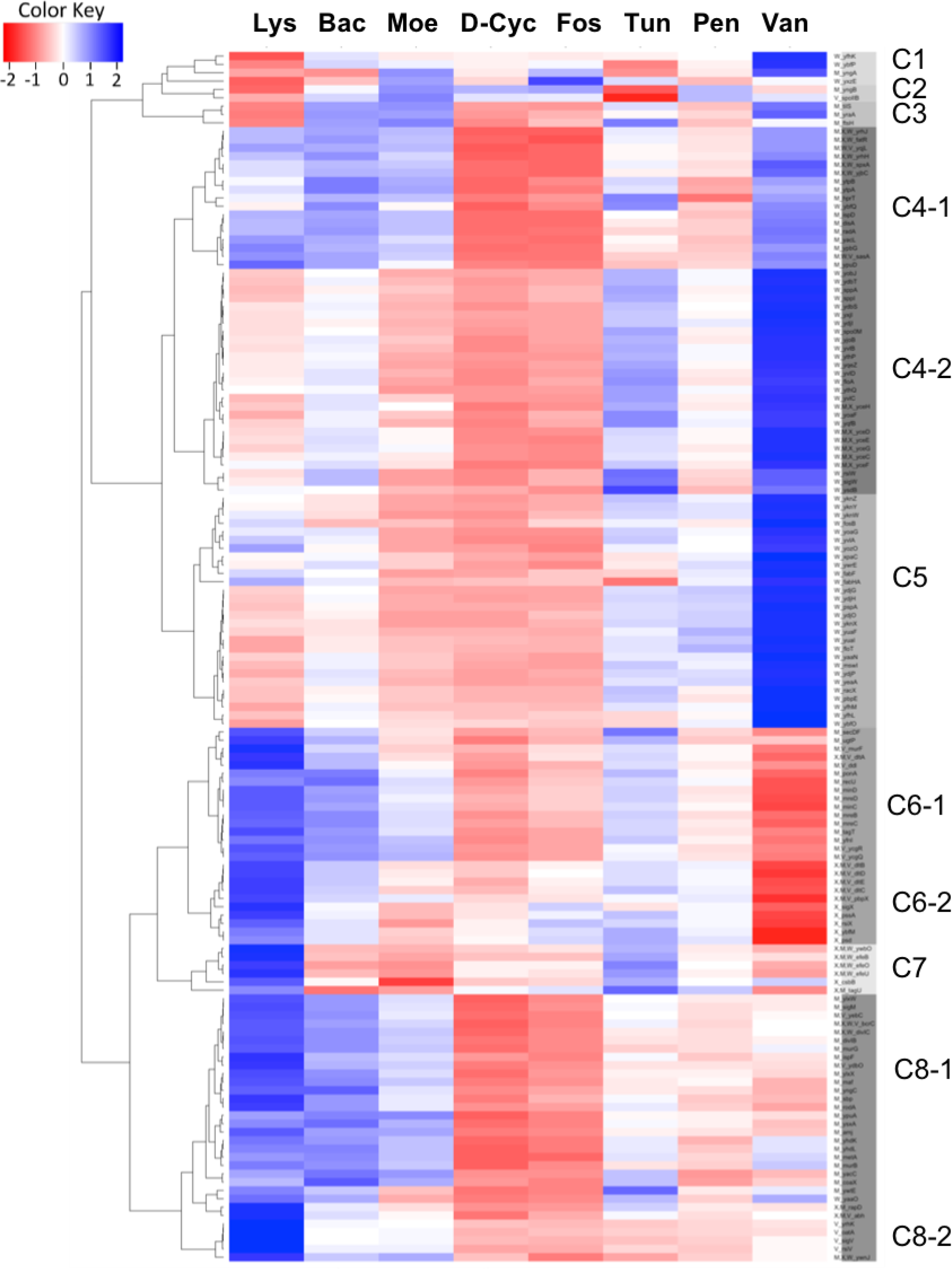
Clustering analysis of the ECF sigma regulons. RNAseq data in log2 was used to generate the heatmap. The RNAseq data applied here is the same as them in Table 1 in the main text which is shown with the data in fold change. The complete data for each gene of the ECF sigma regulons in each treatment conditions and their cluster assignment, shown as C1, C2 et al. at the side of the heatmap, is given in the table attached here.

**Fig. S5.**
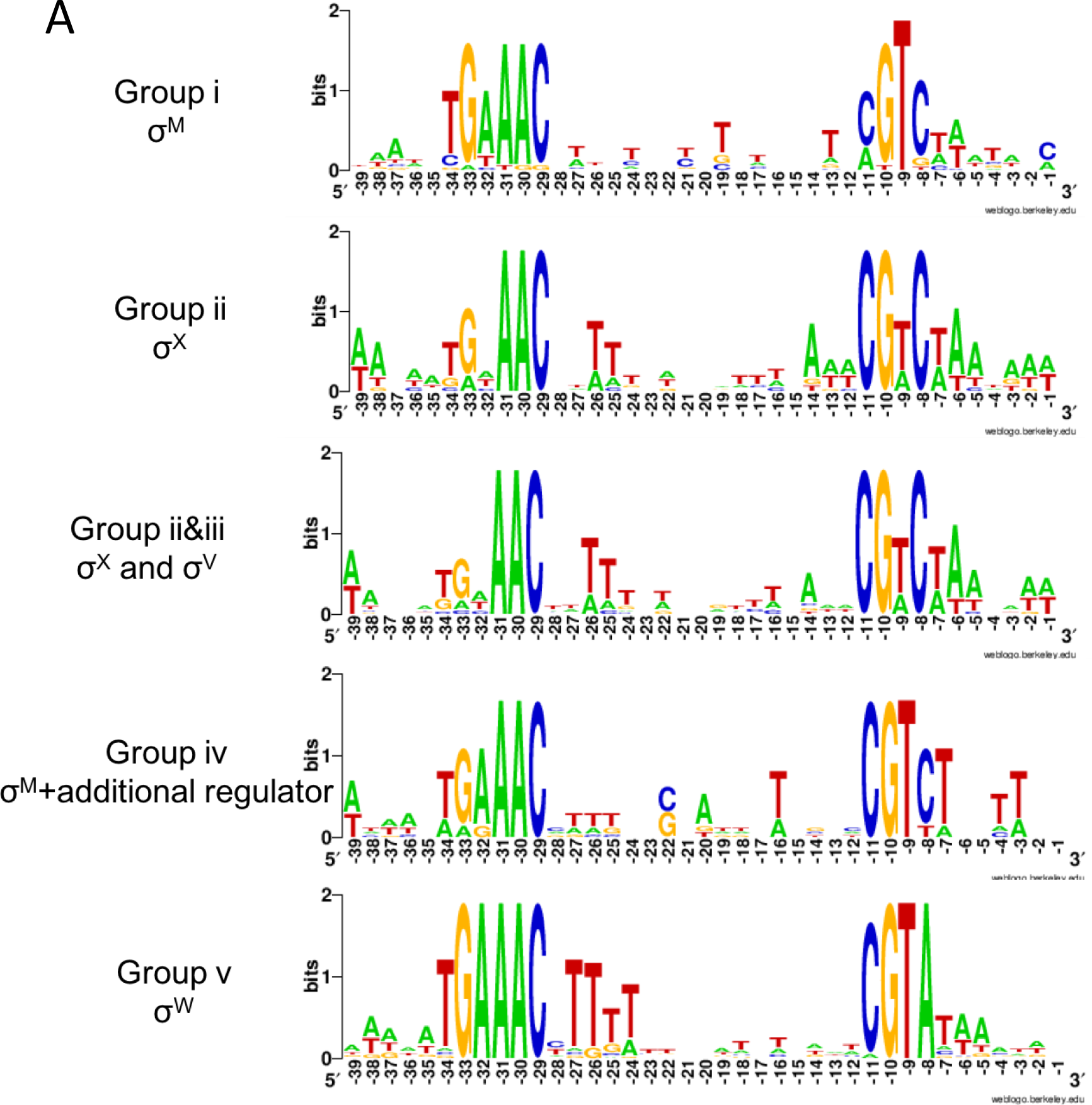
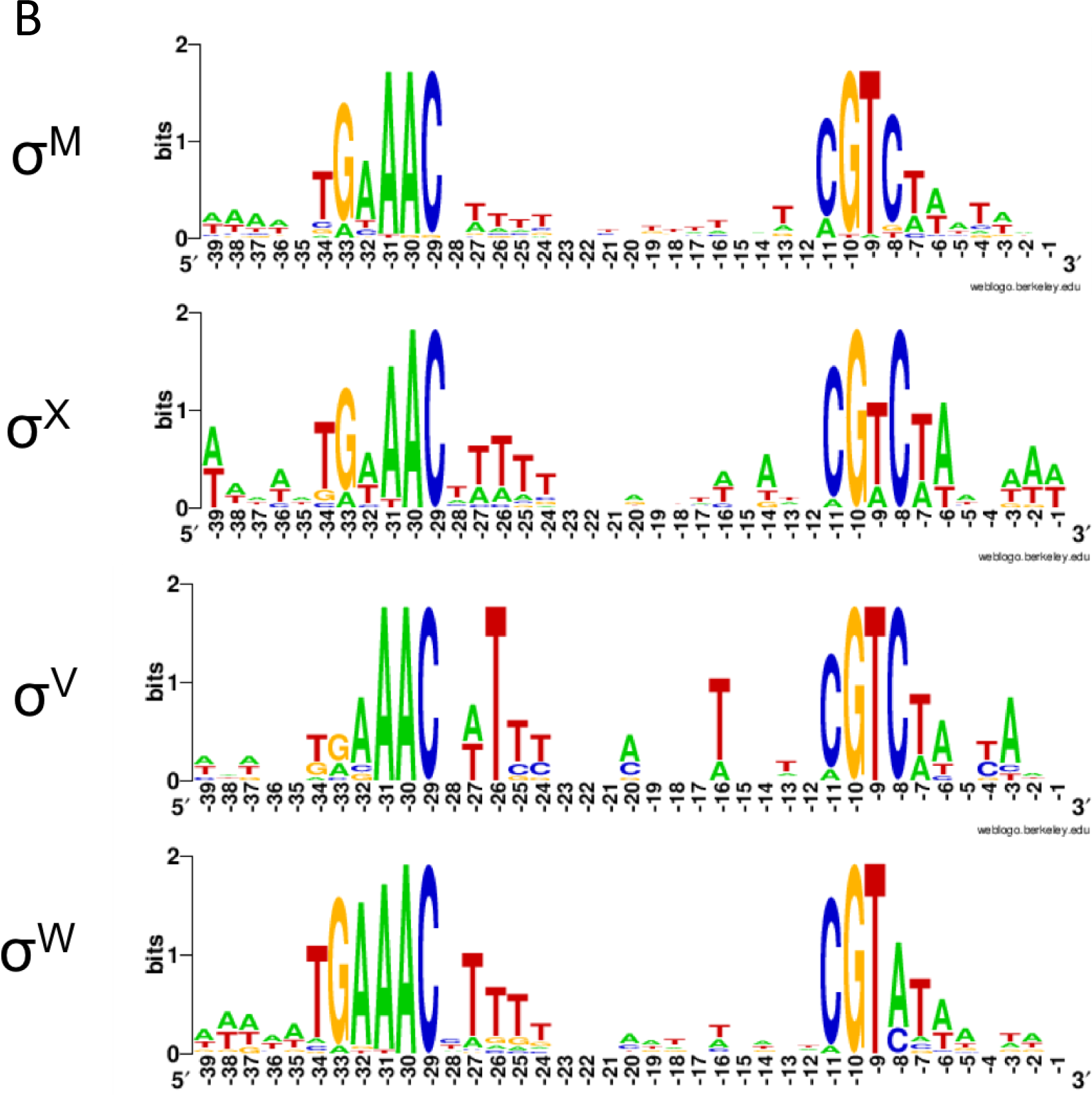
Promoter consensus sequences. (A) represents the promotor consensus sequences for the genes in different groups as shown in Table 2. (B) represents the promotor consensus sequences of the genes previously assigned to each of the four ECF sigma factors σ^W^, σ^M^, σ^X^, and σ^V^.Promoters recognized by ECF σ factors are characterized by conserved sequences near the -35 and -10 regions relative to the transcriptional start site. The promoter sequences and the sources thereof have been shown in Table 2. The sequence logos are generated using WebLogo (http://weblogo.berkeley.edu/).

**Fig. S6.**
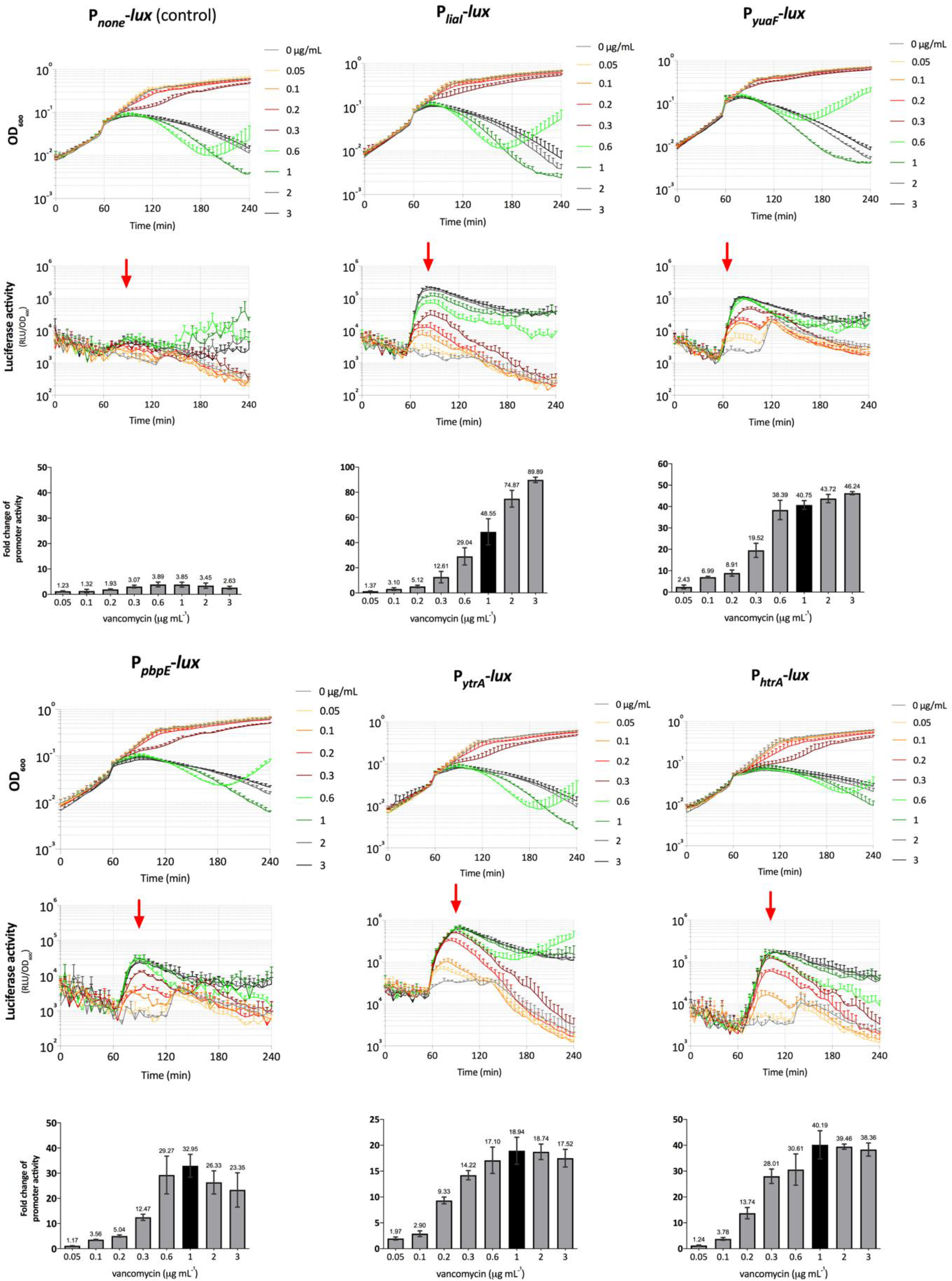
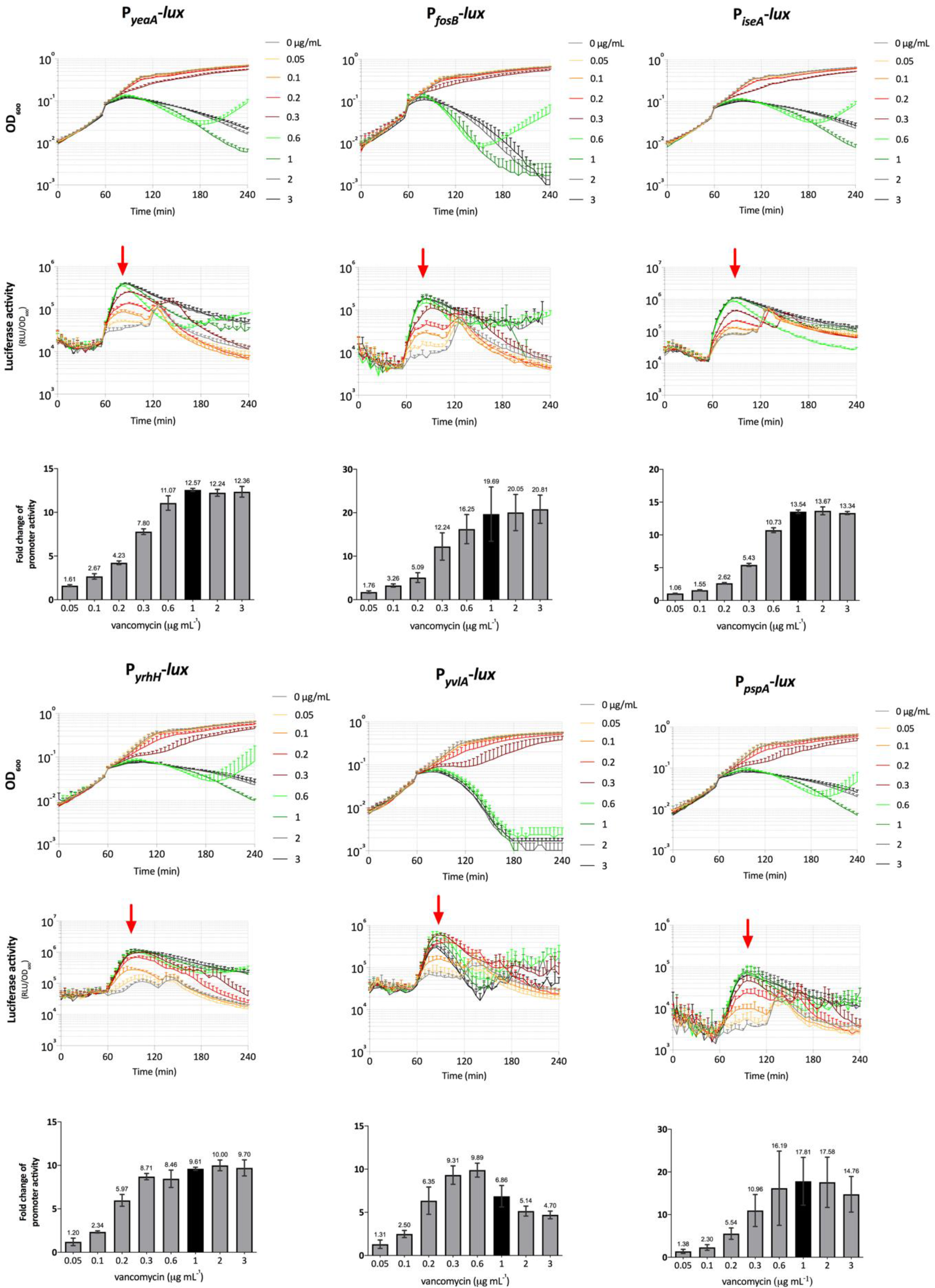
Dose-response of *B. subtilis* biosensors to vancomycin.

**Fig. S7.**
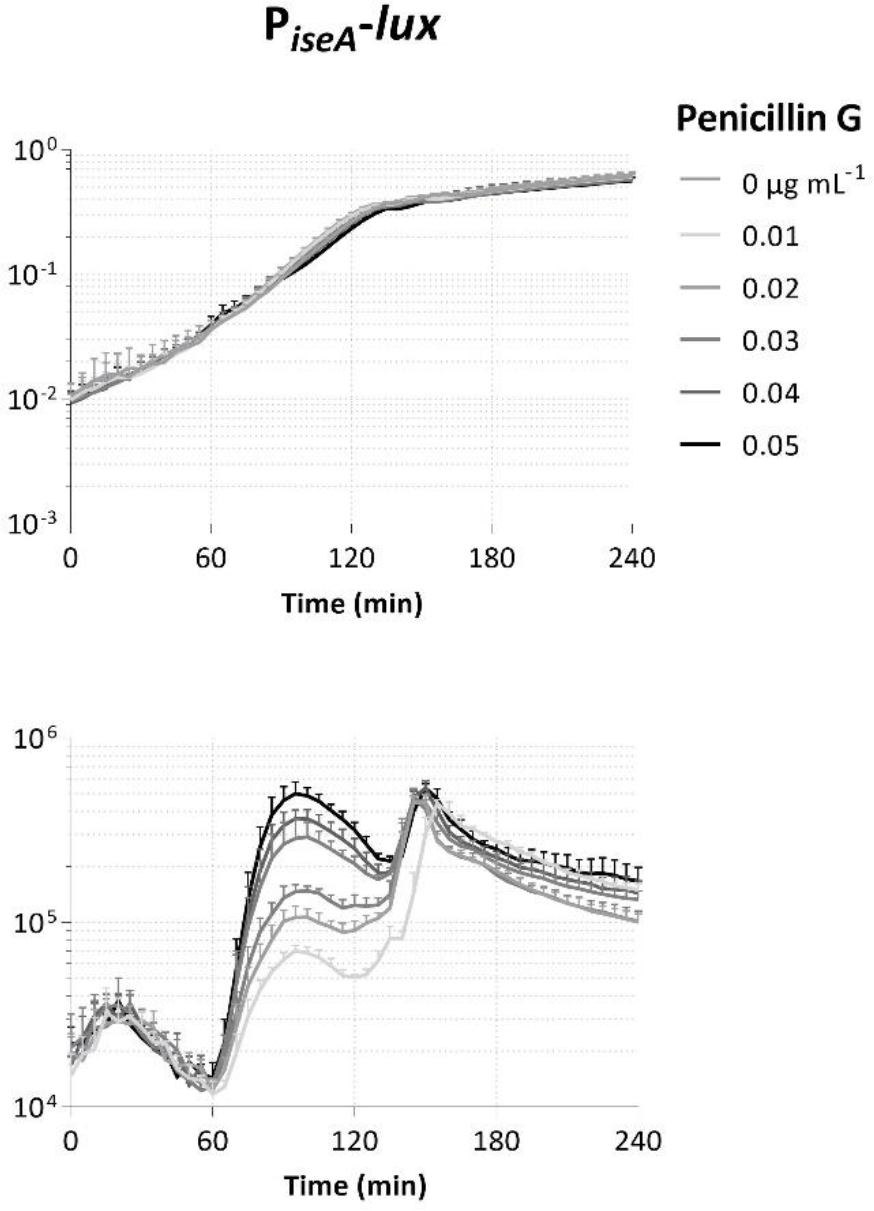
Dose-response of *B. subtilis* biosensor with promoter P*iseA* to penicillin G.

**Fig. S8.**
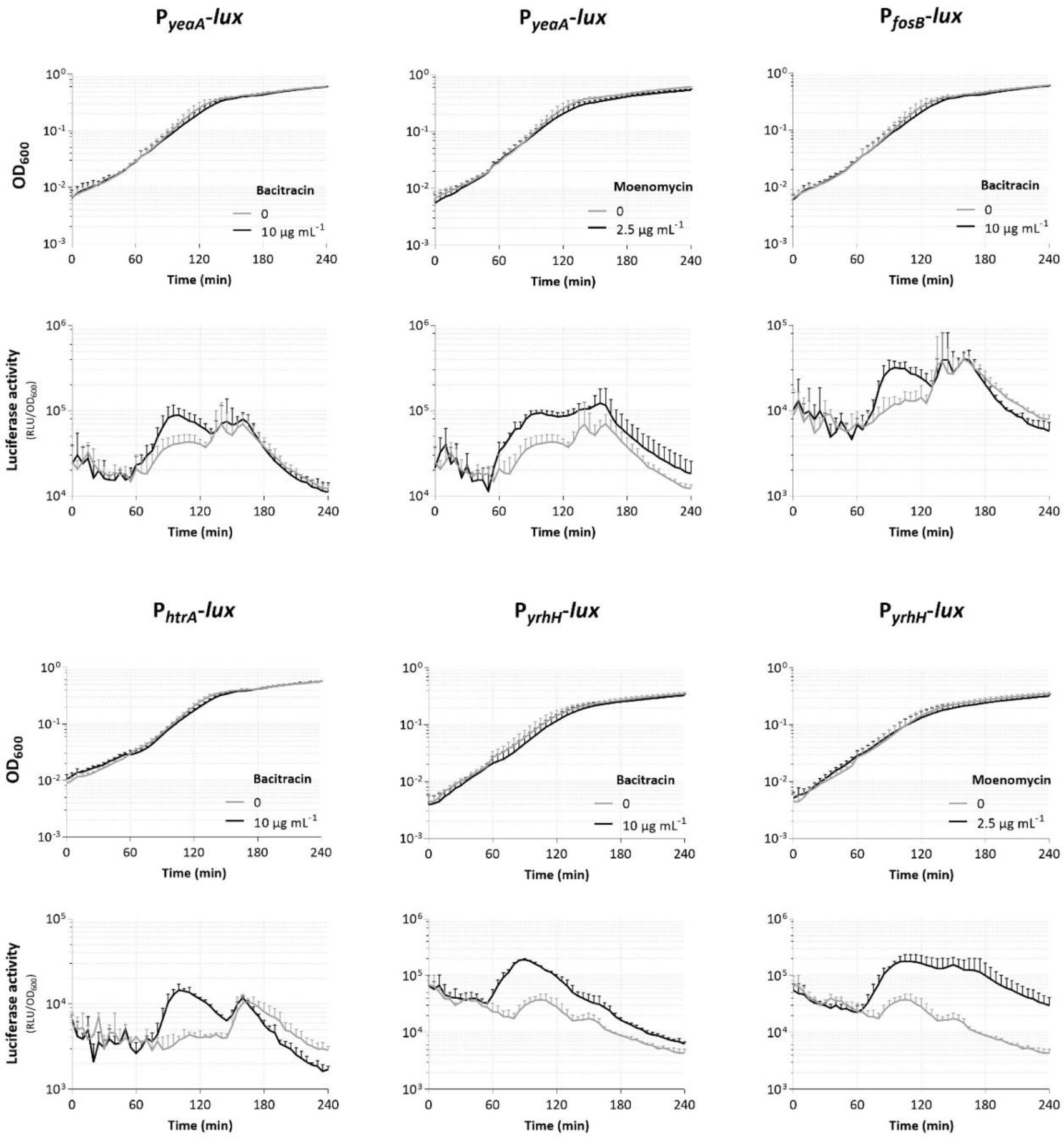
Induction of *B. subtilis* biosensors to bacitracin and/or moenomycin.

